# Single cell transcriptomics, mega-phylogeny and the genetic basis of morphological innovations in Rhizaria

**DOI:** 10.1101/064030

**Authors:** Anders K. Krabberød, Russell J. S. Orr, Jon Bråte, Tom Kristensen, Kjell R. Bjørklund, Kamran Shalchian-Tabrizi

## Abstract

The innovation of the eukaryote cytoskeleton enabled phagocytosis, intracellular transport and cytokinesis, and is responsible for diverse eukaryotic morphologies. Still, the relationship between phenotypic innovations in the cytoskeleton and their underlying genotype is poorly understood. To explore the genetic mechanism of morphological evolution of the eukaryotic cytoskeleton we provide the first single cell transcriptomes from uncultivable, free-living unicellular eukaryotes: the radiolarian species *Lithomelissa setosa* and *Sticholonche zanclea*. Analysis of the genetic components of the cytoskeleton and mapping of the evolution of these to a revised phylogeny of Rhizaria reveals lineage-specific gene duplications and neo-functionalization of α and β tubulin in Retaria, actin in Retaria and Endomyxa, and Arp2/3 complex genes in Chlorarachniophyta. We show how genetic innovations have shaped cytoskeletal structures in Rhizaria, and how single cell transcriptomics can be applied for resolving deep phylogenies and studying gene evolution of uncultivable protist species.

## Introduction

One of the major eukaryotic innovations is the cytoskeleton, consisting of microtubules, actin filaments, actin-related proteins, intermediate filaments and motor proteins. Together these structures regulate the internal milieu of the cell, aid in movement, cytokinesis, phagocytosis, and predation (Dustin 1984, Grain 1986, Vale 2003, Wickstead & Gull 2011, Katz 2012, Cavalier-Smith et al. 2014). Of essential importance, and the main focus of this work, the cytoskeleton of unicellular eukaryotes determines the morphological patterning of the cell.

The evolution of the eukaryotic cytoskeleton is an intriguing story of gene evolution. Homologs to actin and tubulin genes can be found in prokaryotes and Archaea, but the origin of the motor proteins is unclear, lacking distinct homologs in prokaryotes (Vale 2003, Wickstead & Gull 2011). All three major types of motor proteins – kinesin, dynein and myosin – are present in all eukaryote supergroups. Therefore, it is likely that they were already present in the last eukaryotic common ancestor (LECA). Early in the evolution of eukaryotes the cytoskeletal filaments of prokaryotes were given new functions and new motor proteins were invented in addition to a large repertoire of molecules that modify and interact with both the cytoskeleton and the motor proteins (Goldstein 2001, Karcher et al. 2002, Schliwa & Woehlke 2003, Vale 2003, Seabra & Coudrier 2004, Wickstead & Gull 2011).

Most of what we know about the eukaryotic cytoskeleton comes from studies of humans, plants and fungi (Jékely 2007, Wickstead & Gull 2011), but less is known about the genetic machinery and the molecular architecture of the cytoskeleton in non-model single celled eukaryotes (protists). Our current knowledge about the evolution of cytoskeletal genes in protists stems from human pathogens, e.g. *Plasmodium, Toxoplasma* and *Cryptosporidium* (Wickstead & Gull 2011, Burki & Keeling 2014), but virtually nothing is known about how the evolution of these genes has shaped cytoskeletal morphology in other protists.

In this paper we therefore trace the evolution of key cytoskeletal genes in a major group of eukaryotes, Rhizaria, consisting predominantly of understudied single celled protists (Burki & Keeling 2014). Rhizaria is a huge eukaryotic group and harbours species displaying a stunning variety of morphological traits, from naked amoebas to species with delicate and spectacular tests or skeletons. Rhizaria as a group was originally established based on molecular phylogenies that placed the three clades Cercozoa, Radiolaria, and Foraminifera together (Cavalier-Smith 2002, Nikolaev et al. 2004). Although no clearly defined phenotypic synapomorphies for Rhizaria have been described (Pawlowski 2008), there is a common theme to many rhizarians: well-developed pseudopodia which are often reticulose or filose. The different groups of rhizarians use their pseudopodia in different ways: Some form complicated reticulose networks, e.g. many chlorarachniophytes, others use pseudopodia stiffened by microtubules to capture prey, e.g. Radiolaria, to move molecules and organelles, e.g. Foraminifera, or even as oars in Taxopodida (Cachon et al. 1977, Anderson 1978, Sugiyama et al. 2008, Bass et al. 2009). But how this widely different application of pseudopodia has evolved and how the morphological evolution is reflected in changes to cytoskeletal genes is unknown. The cytoskeleton and motor proteins are an integral part of pseudopod development and usage. In the formation of pseudopods in mouse melanoma, actin and myosin interacts in order to make a protrusion in the plasma membrane creating the leading edge of the pseudopod. Nucleators anchor actin to the cell membrane and actin-related proteins (i.e. the Arp2/3-complex) recruits additional actin filaments to form the branching network that supports the pseudopod (Giannone et al. 2007, Mogilner & Keren 2009). The same mechanism drives pseudopod growth and development in the amoeba *Dictyostelium* (Ura et al. 2012). More rigid pseudopods are made with bundles of microtubules that stiffen and support the pseudopods (called axopodia in Radiolaria and reticulopodia in Foraminifera ; Anderson 1983, Lee & Anderson 1991). The microtubules are typically hollow tubes or helical filament composed of alternating α- and β-tubulin subunits (Welnhofer & Travis 1998). The evolution of the molecular components of the cytoskeleton and pseudopodia in Rhizaria, and protists in general, remains unclear.

To understand the evolution of the cytoskeleton and pseudopodia in Rhizaria a fully resolved phylogenetic tree is vital, but getting a stable phylogeny for the entire group has proven problematic. The main issues have been the relationship between Radiolaria and Foraminifera (i.e. Retaria), the monophyly of Cercozoa, as well as the relationship between Rhizaria and its immediate neighbours in the SAR supergroup, Stramenopiles and Alveolata (Burki et al. 2007, 2013, 2016, Parfrey et al. 2010, Krabberød et al. 2011, Sierra et al. 2013, 2015, Katz & Grant 2014, Cavalier-Smith et al. 2015).

Reconstruction of multi-gene phylogenies have been hindered by lacking molecular data from key rhizarian groups (Burki & Keeling 2014). The main reason for scarce data from Rhizaria is lacking knowledge of how to hold species in culture. We have previously applied single cell genomics (combined with gene-targeted PCR) to study the diversity of Retaria, but this method is not optimal to obtain large numbers of protein coding genes as it also covers intergenic regions (Bråte 2012, Krabberød 2011). Other studies targeting the retarian transcriptome have required pooling many, sometimes several hundred cells (Sierra et al. 2013, Balzano et al. 2015), a method not optimal when morphological markers for species identification are missing or hard to define and cultures cannot be established.

Here, we use single cell transcriptomics on two key Rhizaria species (*Sticholonche zanclea* and *Lithomelissa setosa*) to build multi-gene phylogenies and investigate the genetic basis of cytoskeletal differences in Rhizaria. Our aim is to reveal processes at the genetic level that may have caused phenotypic changes to the cytoskeleton, and thereby better understand major morphological transitions in this group of organisms. A key aspect is to understand if gene changes are due to co-option processes, where deeply diverging homologs of cytoskeleton genes have been recruited to new functions, or if novelties of morphologies are caused by innovations of new gene families through gene duplication and neofunctionalization. Are genetic changes specific for each subgroup of Rhizaria or are they common between lineages? And can these changes be used to define homological structures and thereby define morphologically distinct categories of organismal lineages in Rhizaria? To address these questions we apply single cell transcriptomics on two free-living radiolarian species.

## Results

### Single cell transcriptomics of two uncultured protists

We generated cDNA libraries from two radiolarian specimens: *Lithomelissa setosa* and *Sticholonche zanclea* (Figure 1). The cDNA was sequenced on the Illumina MiSeq platform, 300bp paired end. This resulted in 19,894,654 reads for *S. zanclea* and 11,590,658 for *L. setosa*, which were *de novo* assembled using the Trinity platform (Haas et al. 2013). Assembly resulted in two Single Cell Transcriptomes (SCT) with 4,749 predicted genes for *S. zanclea and* 2,122 predicted genes for *L. setosa* (Table 1). Subsampling and re-assembly of reads showed that the sequencing threshold for both libraries was close to maximum (Figure 1-figure supplement 1). We assessed the suitability of the data for phylogenomic reconstruction by using the BIR pipeline for single gene alignment and tree construction (Kumar et al. 2015). Using 255 seed alignments covering the eukaryote Tree of Life (Burki et al., 2012) we identified 54 and 16 corresponding orthologous gene sequences from *S. zanclea* and *L. setosa* respectively. In addition BIR extracted 3,534 gene sequences from Marine Microbial Eukaryote Transcriptome Sequencing Project, MMETSP (Keeling et al., 2014) and 793 proteins from GenBank with TaxID 543769 (Rhizaria) and added these to their corresponding alignments (See supplementary table S1). After concatenation of all gene alignments we had a super-matrix consisting of 91 taxa and 54,898 amino acids (255 genes).

**Figure 1.**
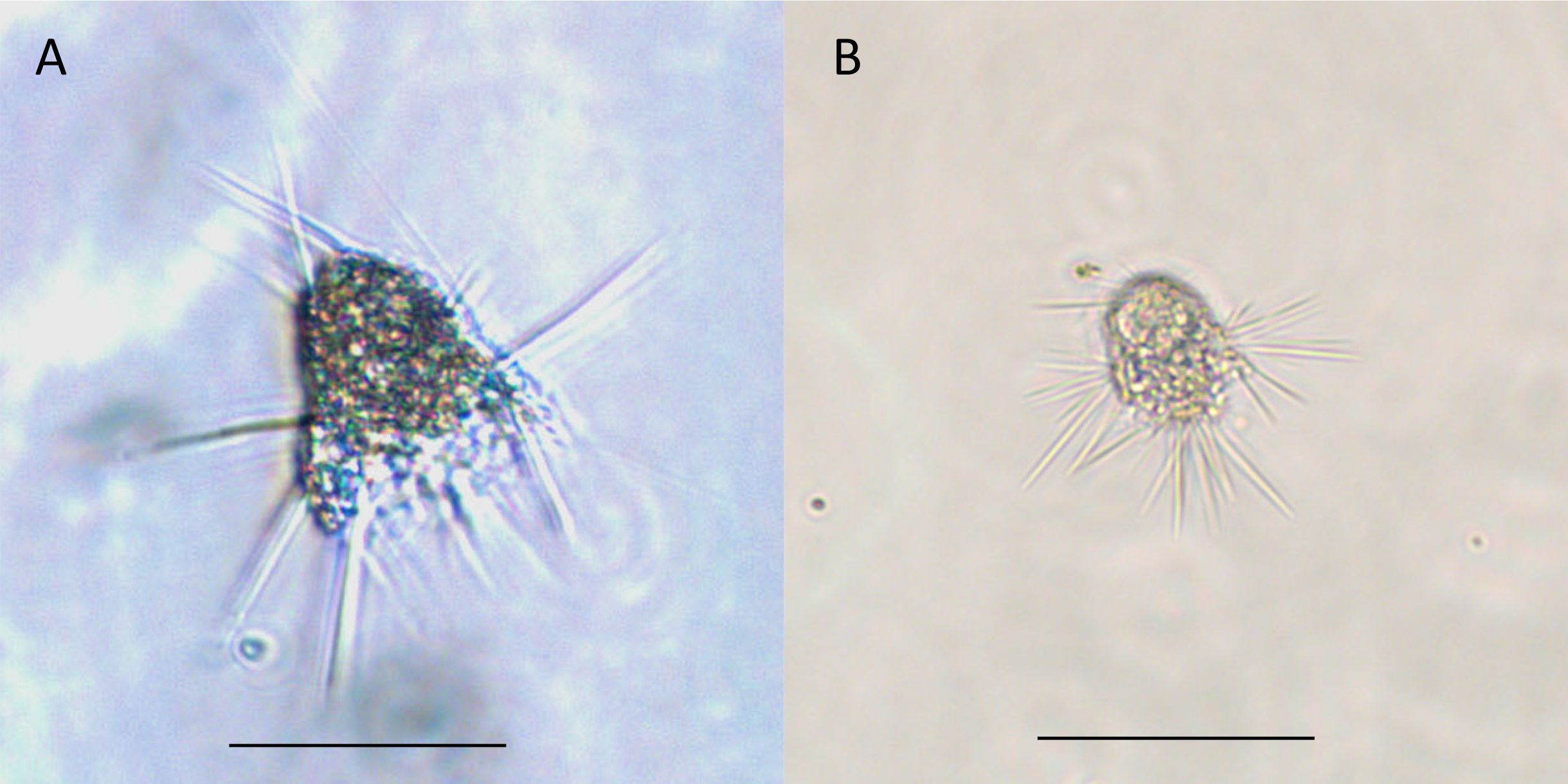
The two specimens sequenced. A) *Lithomelissa setosa*, B) *Sticholonche zanclea*. Scale bar 50 μm.

**Table 1:** Single cell transcriptome statistics

| Species name | Raw Reads | Contigs<sup>1</sup> | GC-content (%) | Predicted Genes<sup>2</sup> |
| --- | --- | --- | --- | --- |
| *Sticholonche zanclea* | 19,894,654 | 19,509 | 53.5 | 4,749 |
| *Lithomelissa setosa* | 11,590,658 | 12,212 | 48.8 | 2,122 |

<sup>1</sup> Number of contigs assembled by Trinity (Haas et al. 2013). <sup>2</sup>The number of genes predicted by TransDecoder in the Trinity platform.

**Table 2:** Rhizarian transcriptomes from MMETSP (Keeling et al. 2014) used in this study.

| Sample Id | Phylum | Species | Strain | Transcripts<sup>1</sup> |
| --- | --- | --- | --- | --- |
| MMETSP0040 | Chlorarachniophyta | *Lotharella oceanica* | CCMP622 | 17,354 |
| MMETSP0041 | Chlorarachniophyta | *Lotharella globosa* | LEX01 | 25,644 |
| MMETSP0042 | Chlorarachniophyta | *Amorphochlora amoebiformis* * | CCMP2058 | 23,387 |
| MMETSP0045 | Chlorarachniophyta | *Bigelowiella natans* | CCMP 2755 | 22,651 |
| MMETSP0109 | Chlorarachniophyta | *Chlorarachnion reptans* | CCCM449 | 26,481 |
| MMETSP0110 | Chlorarachniophyta | *Gymnochlora* sp. | CCMP2014 | 15,507 |
| MMETSP0111 | Chlorarachniophyta | *Lotharella globosa* | CCCM811 | 19,670 |
| MMETSP0112 | Chlorarachniophyta | *Lotharella globosa* | CCCM811 | 11,910 |
| MMETSP0113 | Chlorarachniophyta | *Norrisiella sphaerica* | BC52 | 14,550 |
| MMETSP0186 | Cercozoa | *Minchinia chitonis* | Missing | 461 |
| MMETSP1052 | Chlorarachniophyta | *Bigelowiella natans* | CCMP623 | 24,186 |
| MMETSP1318 | Chlorarachniophyta | *Partenskyella glossopodia* | RCC365 | 15,025 |
| MMETSP1358 | Chlorarachniophyta | *Bigelowiella natans* | CCMP1242 | 18,273 |
| MMETSP1359 | Chlorarachniophyta | *Bigelowiella longifila* | CCMP242 | 15,959 |
| MMETSP1384 | Foraminifera | *Ammonia* sp. | Missing | 31,225 |
| MMETSP1385 | Foraminifera | *Elphidium margaritaceum* | Missing | 25,184 |

<sup>1</sup>The number of amino acid sequences for the sample. \* *Amorphochlora amoebiformis* is called *Lotharella* *amoebiformis* in MMETSP, but it was moved to the genus *Amorphochlora* by Ishida et al., (2011)

### Bayesian GTRCAT trees show consistent phylogeny for SAR and subgroups

In the Bayesian analysis of the full dataset using the CATGTR model (255 genes 54,898 AA, 91 taxa, Figure 2), Stramenopiles and Alveolates formed a clade, with Rhizaria as sister; branches for these groups are fully supported (1.00 posterior probability (pp)). Haptophytes appeared as sister to SAR (0.87 pp). The relationship and support values did not change for SAR with the removal of fast evolving sites (Figure 2-figure supplement 1; supplementary table S4). The haptophytes, jumped to a position basal to Archaeplastida (0.82 pp) when four bins of fast evolving sites were removed (supplementary table S4).

**Figure 2.**
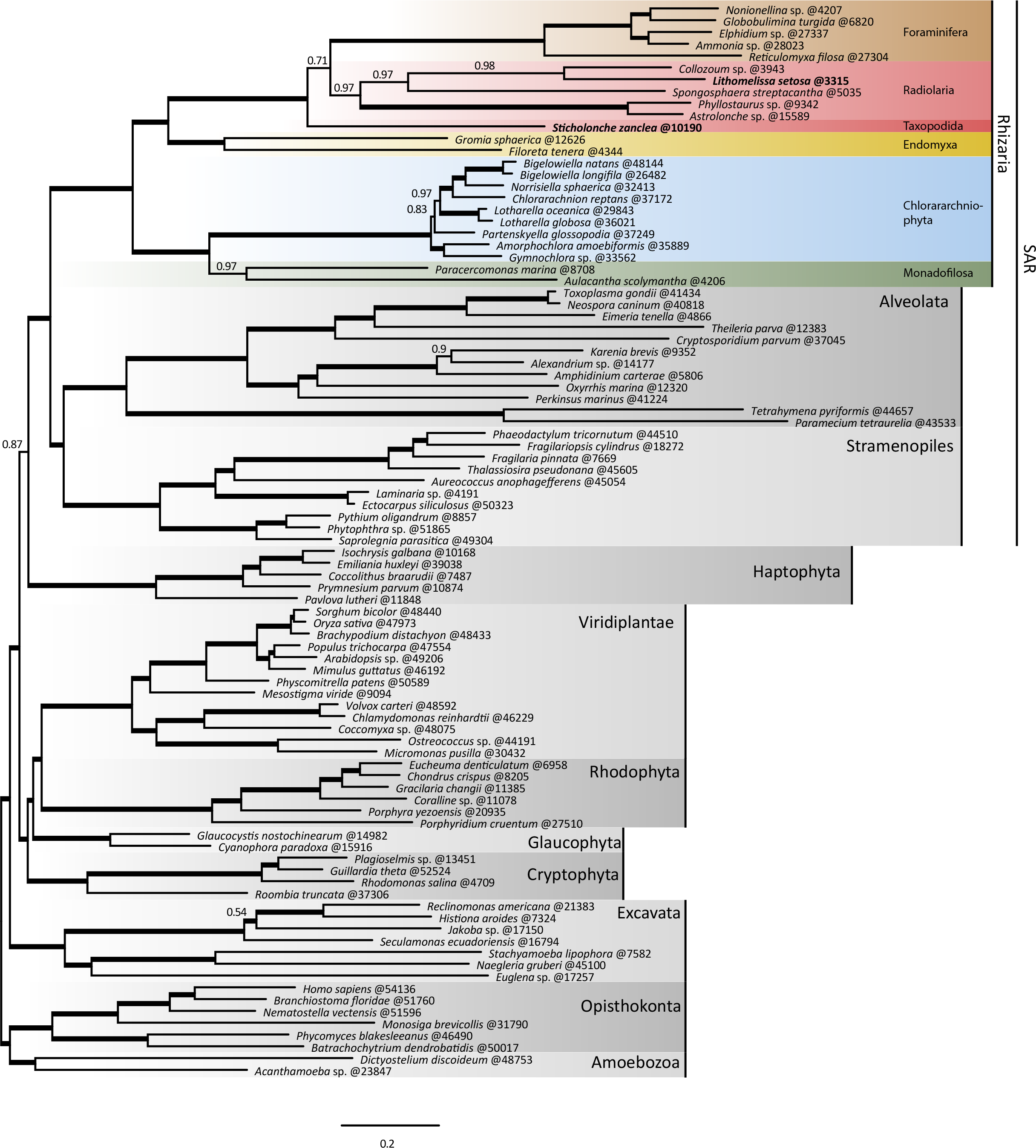
Bayesian phylogeny with the CATGTR model, 255 genes, 54,898 AA, and 91 taxa, maxdiff 0.2666. Species sequenced for this paper in bold. Thick branches represent maximal support (posterior probability = 1). Number after ‘@’ is concatenated sequence length. Important clades in Rhizaria are coloured for easier identification: brown = Foraminifera, dark red = Taxopodida, red = Radiolaria, yellow = Endomyxa, blue= Chlorarachniophyta (Filosa), and green = Mondaofilosa (Filosa). The scale bar equals the mean number of substitutions per site. The changing support values for selected branches depending on the number of genes, and the number of fast evolving sites removed are shown in Figure 2-figure supplement 1.

Within Rhizaria, the three groups Foraminifera, Radiolaria and Taxopodida, all monophyletic, formed a cluster (i.e. Retaria) with maximum support even when fast evolving sites were removed (i.e. always 1.00 pp). Radiolaria and Foraminifera were placed together as a monophyletic group (0.71 pp) with *S. zanclea* branching off as sister to them both. This topology remained constant after removing fast evolving sites (Figure 2 – figure supplement 1; supplementary table S4). The posterior probability for the monophyly of Radiolaria together with Foraminifera, i.e. excluding *S. zanclea*, increased to 0.97 when fast evolving sites were removed (Figure 2 – figure supplement 1). Endomyxa was monophyletic (1.00 pp) and always sister to Retaria with full support (1.00 pp), rendering Cercozoa paraphyletic. Filosa was monophyletic in all analyses (1.00 pp).

**Figure 3.**
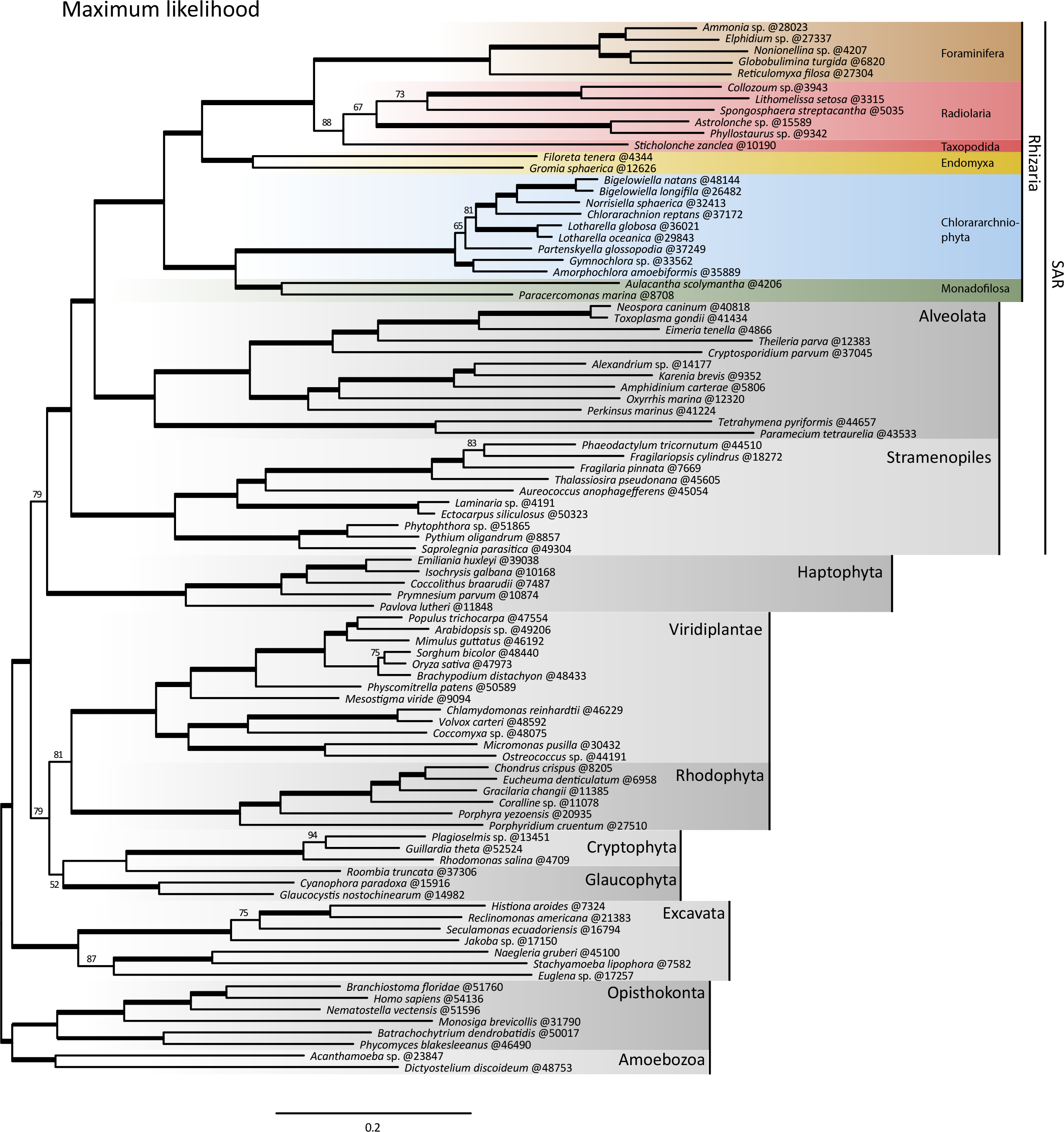
Maximum likelihood with the LG model, 255 genes, 54,898 AA, and 91 taxa. Species sequenced for this paper in bold. Thick branches represent maximal support (bootstrap = 100 %). Number after ‘@’ is concatenated sequence length. As in figure 2 important clades in Rhizaria are coloured for easier identification: brow = Foraminifera, dark red = Taxopodida, red=Radiolaria, yellow = Endomyxa, blue= Chlorarachniophyta (Filosa), and green = Mondaofilosa (Filosa). The scale bar equals the mean number of substitutions per site. The changing support values for selected branches depending on the number of genes, and the number of fast evolving sites removed are shown in Figure 3 – figure supplement 1.

### ML trees converged towards Bayesian topology after removal of fast evolving sites

In contrast, the maximum likelihood (ML) analysis of the full dataset using the LG model (255 genes, 54,898 AA, 91 taxa, Figure 3), grouped Alveolata with Rhizaria instead of the Stramenopiles (96% bootstrap support (bs)). Retaria was recovered with high support (88% bs) as in the Bayesian tree, but *S. zanclea* was no longer placed ancestrally to radiolarians and Foraminifera. Instead *S. zanclea* was sister to Radiolarians (88% bs). Importantly, however, *S. zanclea* changed to a basal position in Retaria after removal of fast evolving sites, consistent with all the GTRCAT Bayesian trees (Figure 3 – figure supplement 1; supplementary table S4).

Removal of fast evolving sites did not change the monophyly of Foraminifera and Radiolaria (excluding *S. zanclea*) or the sister relation between alveolates and Rhizaria, but the support values were reduced in both instances to 50% bs for Radiolaria together with Foraminifera and 67% bs for the alveolates together with Rhizaria (Figure 3 – figure supplement 1; supplementary table S4). Endomyxa and Retaria group together with full support as in the Bayesian analysis (100% bs), making Cercozoa paraphyletic. As in the Bayesian phylogeny haptophytes appeared as sister to SAR (77% bs) and changed position basal to the plants, glaucophytes and cryptomonads (73% bs) after removal of fast evolving sites. Species with more than 10% missing data in the final concatenated data matrix were placed on the ML phylogeny using the Evolutionary Placement Algorithm (Berger & Stamatakis 2011). Five species were placed in Endomyxa, five in Filosa, two in Radiolaria, and finally ten species in Foraminifera (Figure 14).

### Influence of fast evolving sites and the choice of model on the phylogeny

The discrepant topologies of the Bayesian (CATGTR) and ML (LG) trees could be due to the different models implemented in these two approaches. We assessed the influence of these two models by running Bayesian inferences using the LG model (the opposite: running ML with a CATGTR model is currently not possible). This was done on a smaller alignment to reduce the computational burden (146 genes, 33,081 AA, 91 taxa, see methods for further explanation). The resulting Bayesian tree showed an important result: *S. zanclea* now grouped with Radiolaria (0.67 pp, Figure 3 – figure supplement 1) as in the ML (LG) tree, and not as sister to Foraminifera and Radiolaria, as in all Bayesian trees with the CATGTR model. Other branching patterns in the Rhizaria phylogeny were unaffected.

We repeated the ML (LG) analyses after removing fast evolving sites on the full dataset as well as the reduced dataset. While alveolates and Rhizaria formed a clade in the full and small dataset (85% bs, Figure 3- figure supplement 1), removal of four categories of fast evolving site moved alveolates to the stramenopiles in the dataset with 146 genes (74% bs, Figure 3 – figure supplement 1; supplementary table S4), a result congruent with the Bayesian topology. The support for alveolates together with Rhizaria was also weakened in the dataset with 255 gene when four categories of fast evolving sites where removed, from 96% bs, to 67% bs. When Foraminifera was excluded from the 255 gene dataset with four categories removed, Stramenopiles and Alveolata formed a group with Rhizaria as sister (57% bs. Table S4).

### Actin radiation in Rhizaria and unique duplications in Retaria and Endomyxa

We identified 6 actin sequences in our SCTs. From MMETSP we identified 18 foraminiferan actin and 18 chlorarachniophyte sequences. Phylogenetic analysis of these and other available actin sequences retrieved from GenBank and Pfam revealed that Retaria (including *S. zanclea*) have two distinct paralogs of actin – actin1 and 2 – where actin2 is fully supported (Figure 4). Actin1 is supported in the Bayesian analysis (0.87 pp) but not in by ML analysis. Actins from Endomyxa form a weakly supported monophyletic group with retarian actin2 (14% bs/0.68 pp). This clade, in turn, groups together with retarian actin1 (59% bs/ 0.97 pp), and is a synapomorphy for Retaria and Endomyxa. There are possibly three paralogs of endomyxean actin, (named a-c in Figure 4) albeit lacking support (a: 32%bs /0.6 pp, b: 16 %bs /-and c: 22 %bs /0.62 pp, Figure 4).

**Figure 4.**
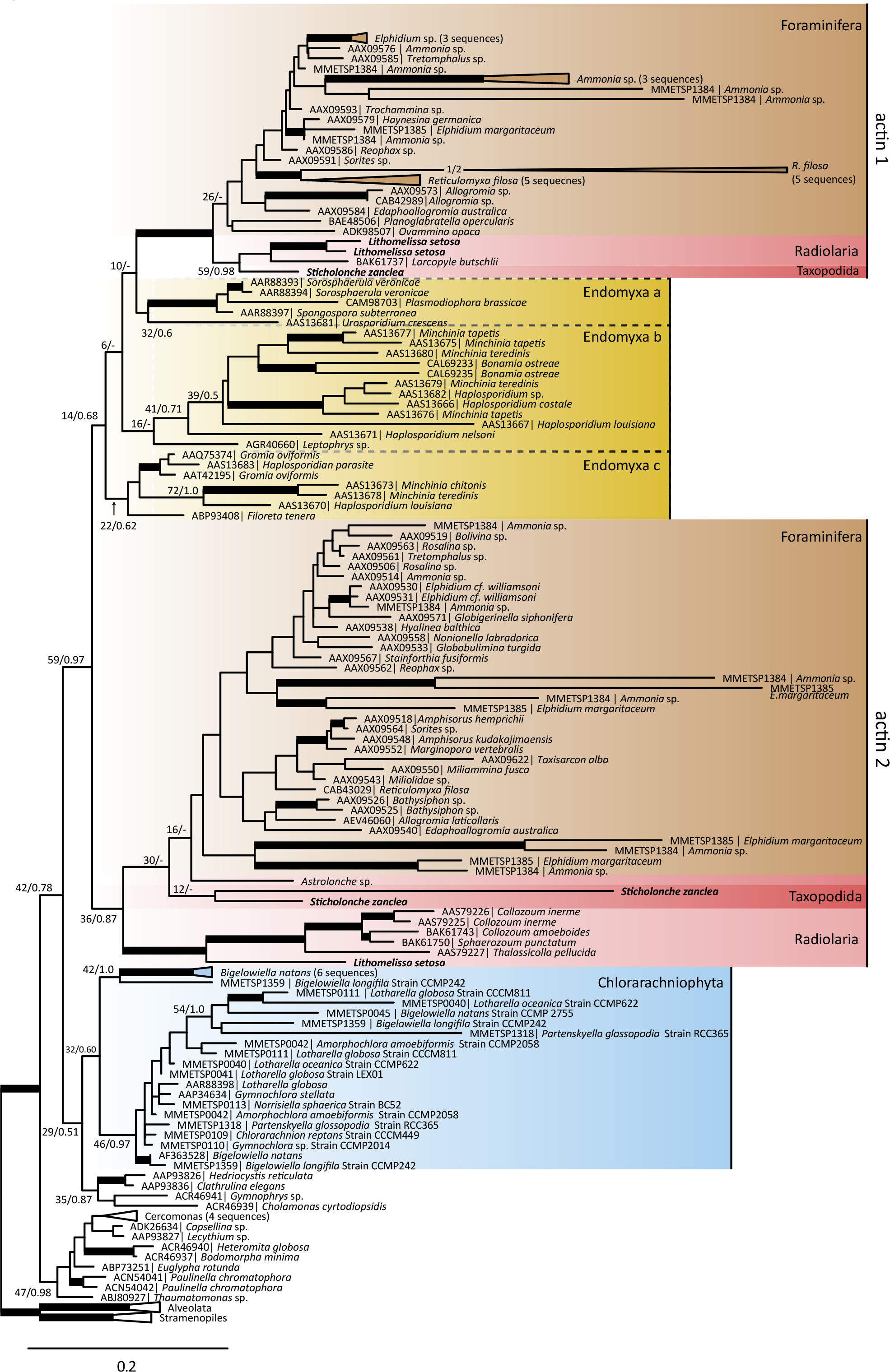
Actin phylogeny (229 taxa, 374 AA). Thick branches represents bootstrap > 75% and posterior probability > 0.9. Some branches are collapsed to save space. Support values for selected nodes discussed in the text added for clarity. The scale bar equals the mean number of substitutions per site. The colouring scheme is the same as in figure 2. (Brown = Foraminifera, red = Radiolaria/Taxopodida, yellow = Endomyxa and blue= Filosa).

### Arp2/3 complex gene duplication in Chlorarachniophyta

Of the seven genes in the Arp2/3 complex, which is responsible for branching of actin filaments and recruitment of new actin, we identified *Arp2, Arp3, ARPC2* and *ARPC5* from *S. zanclea*, but only *Arp2* from *L. setosa* (Figure 5). From MMETSP we identified sequences of all seven genes from both Chlorarachniophyta and Foraminifera. Phylogenetic analysis of these genes revealed that all chlorarachniophytes have two distinct paralogs of both *Arp2* and *APRC1* (Figure 5A), recovered with maximum support (100% bs/1.0 pp). No other species of Rhizaria has undergone the same gene duplication in the Arp2/3 complex as the chlorarachniophytes (Figure 5B).

**Figure 5.**
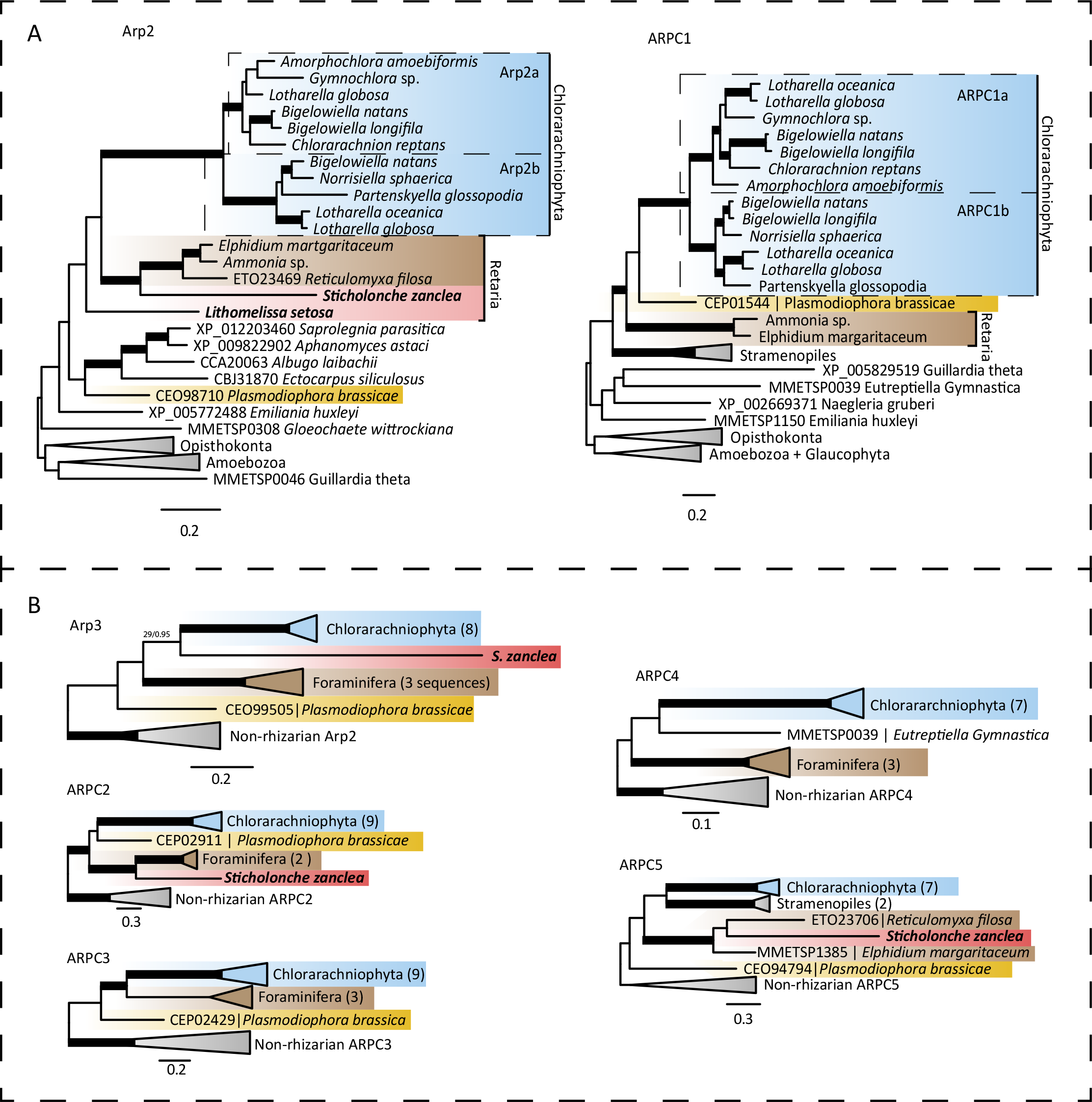
Phylogenies of the seven genes in the Arp2/3 complex. *Arp2* (39 taxa, 373 AA), Arp3 (33 taxa, 403 AA), *ARPC1* (34 taxa, 328 AA), *ARPC2* (24 taxa, 303 AA), *ARPC3* (30 taxa, 181 AA), *ARPC4* (23 taxa, 169 AA) and *ARPC5* (29 taxa, 151 AA). Colouring of groups as in figure 2 (Brown = Foraminifera, red= Radiolaria, yellow = Endomyxa and blue= Filosa). Thick branches represents bootstrap > 75% and posterior probability > 0.9 and the scale bar equals the mean number of substitutions per site. (A) The two genes with a recent duplication in Chlorarachniophyta (*Arp2* and *APRC1*). (B) The five genes without duplication in Chlorarachniophyta (*ARP3, ARPC2, ARPC3, ARPC4, ARPC5*).

### Neofunctionalization of Arp2/3 in Chlorarachniophytes

Comparative evolutionary analyses of the duplicated Arp2/3 complex genes (*Arp2* and *ARPC1*) were performed by examining the evolutionary rates for each paralog, and then mapping the genes to structural models using Consurf (Ashkenazy et al. 2010, Celniker et al. 2013). The analysis showed that the two different forms of Arp2 (Arp2a and Arp2b, Figure 6) and the two different forms of ARPC1 (ARPC1a and ARPC1b, Figure 7) follow a pattern where the most conserved sites are localized inside the protein structure. Comparison of the surface between the two Arp2 proteins (Arp2a and Arp2b, Figure 6) show shared conserved residues in contact surfaces against other proteins in the Arp2/3 complex (colored green in Figure 8). Similarly, the two different paralogs of ARPC1 (ARPC1a and ARPC1b, Figure 8) show shared conserved sites localized inside the complex. In contrast, the surfaces of the two Arp2 and ARPC1 copies show more variable substitution rates, and all paralogs have patches with mutually exclusive conserved residues (figure 8). As the surfaces of Arp2 and ARPC1 are responsible for the recruitment of the daughter filament and are important for anchoring the two actin strands to each other, the divergent substitution patterns imply that the pairs of paralogs have evolved into different directions from the ancestral gene and thereby undergone sub-functionalization.

**Figure 6.**
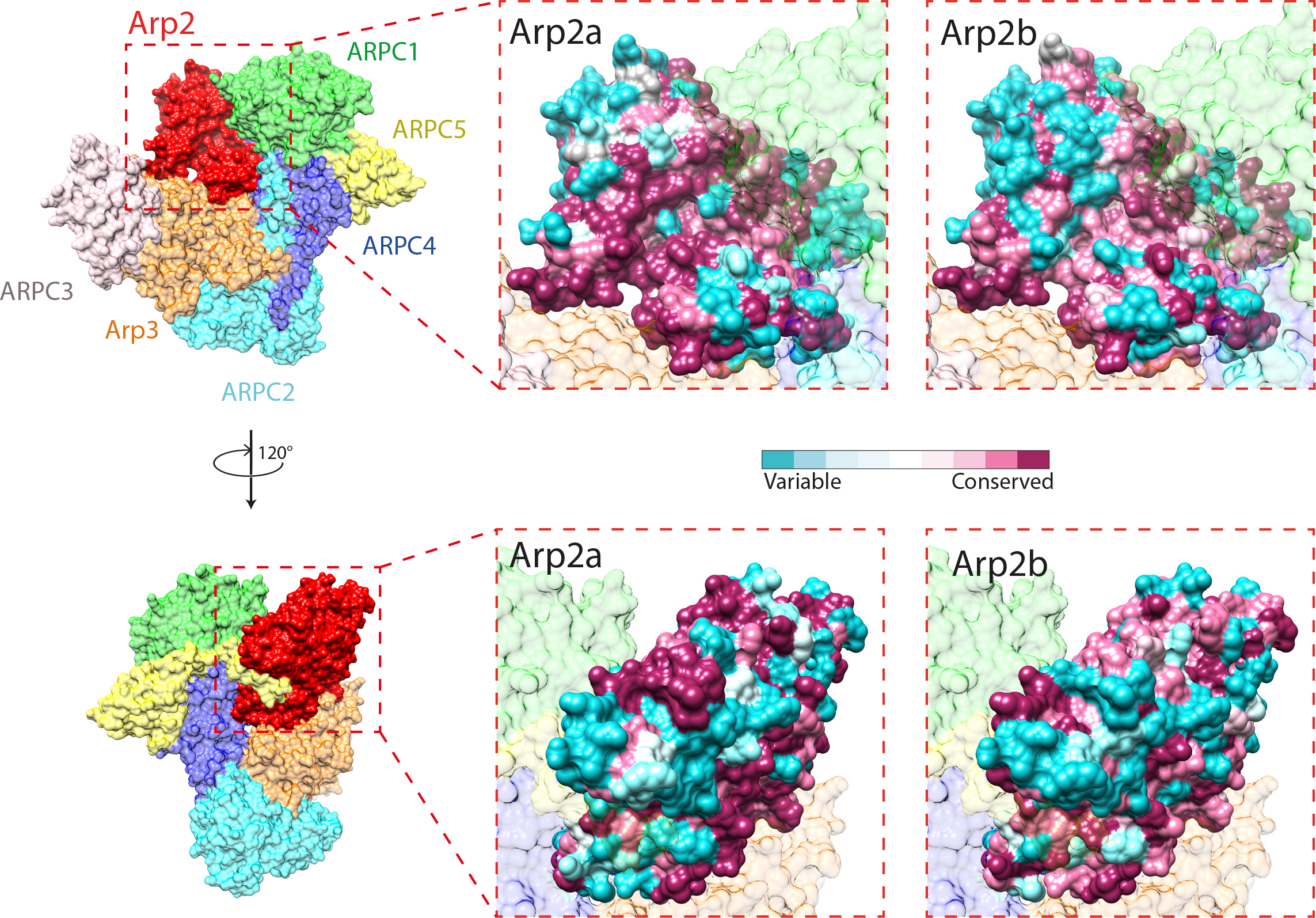
Molecular models of the Arp2a and Arp2b paralogs in Chlorarachniophyta with evolutionary rates from Consurf superimposed on the Arp2/3 complex (PDB accession 4JD2; Arp2 red, Arp3 orange, APRC1 green, ARPC2 cyan, ARPC3 pink, ARPC4 blue, ARPC5 yellow). Residues are coloured according to the evolutionary rates calculated by Consurf. Turquoise residues are highly variable and maroon means conserved residues.

**Figure 7.**
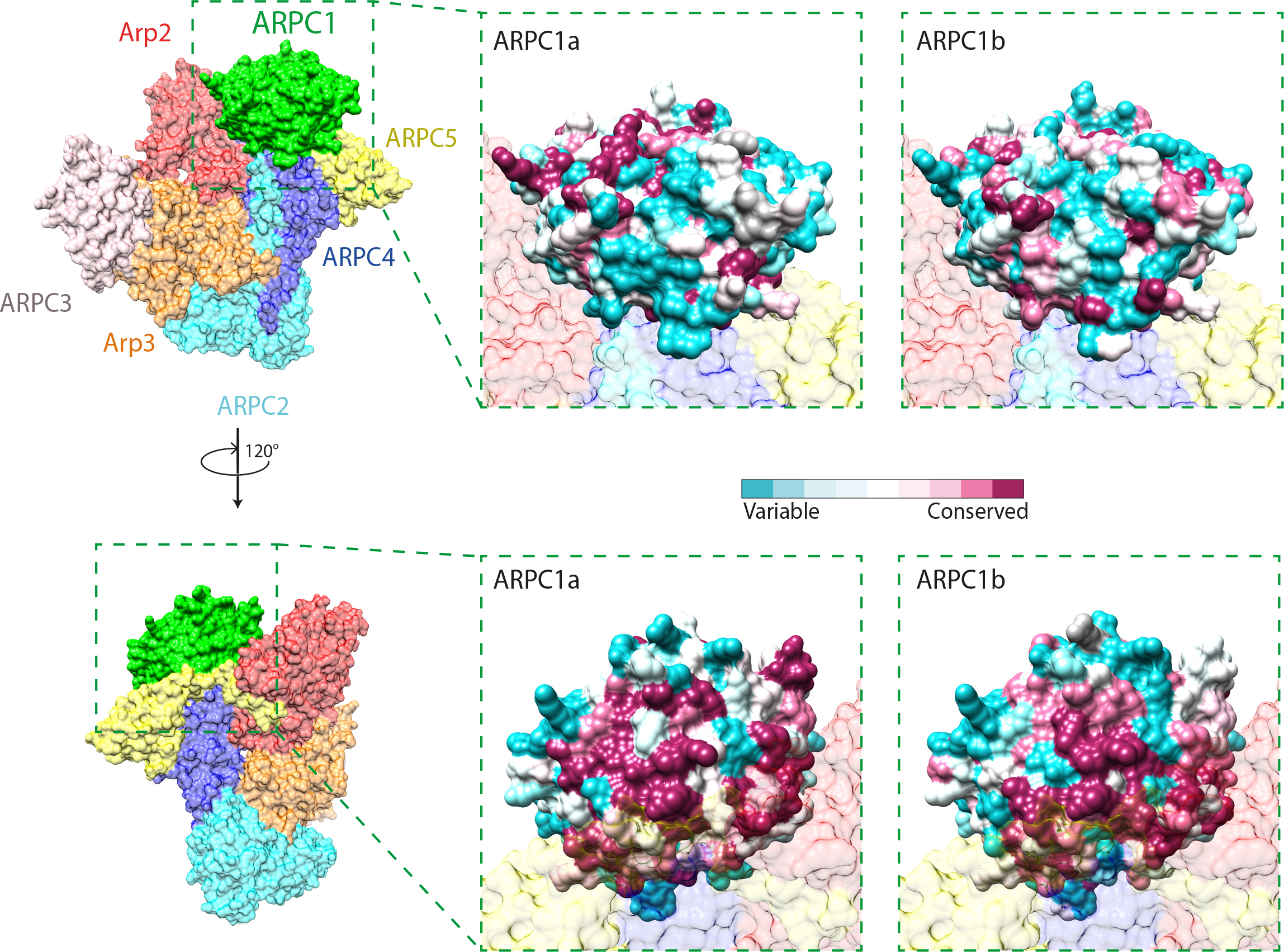
Molecular models of the ARPC1a and ARPC1b paralogs in Chlorarachniophyta with evolutionary rates from Consurf superimposed on the Arp2/3 complex (PDB accession 4JD2; Arp2 red, Arp3 orange, APRC1 green, ARPC2 cyan, ARPC3 pink, ARPC4 blue, ARPC5 yellow). Residues are coloured according to the evolutionary rates calculated by Consurf. Turquoise residues are highly variable and maroon means conserved residues.

**Figure 8.**
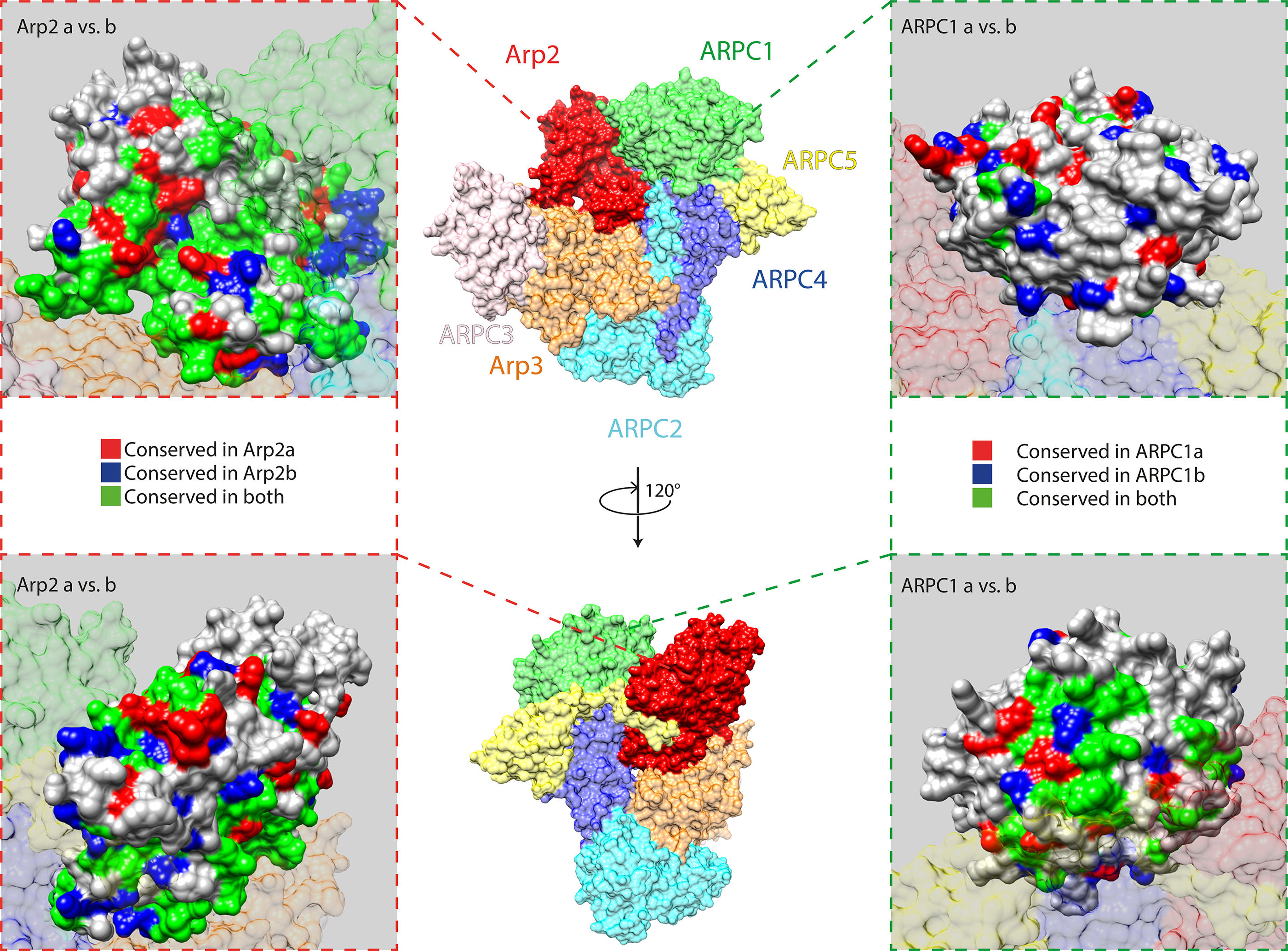
Comparison of conserved residues between the two paralogs of Arp2 and the two paralogs of ARPC1 in Chlorarachniophytes superimposed on PDB accession 4JD2. A) Conserved residues from the two different paralogs of Arp2 (Arp2a and Arp2b). Red represent residues conserved in Arp2a only, blue are conserved in Arp2b only, while green residues are conserved in both paralogs. B) Conserved site from the two different paralogs of ARPC1 (ARPC1 and ARPC1b). Red sites represent residues conserved in ARPC1 only, blue are conserved in ARPC1 only, while green residues are conserved in both paralogs.

### Myosin evolution in Rhizaria

We identified 133 myosin transcripts from MMETSP with rhizarian origin. A phylogenetic reconstruction of the newly identified rhizarian myosins together with already published myosin classes spanning a broad taxonomical distribution of eukaryotes (Richards & Cavalier-Smith 2005, Sebé-Pedrós et al. 2014) revealed the presence of two known classes (If and IV) and three previously unknown classes of myosin in Rhizaria (XXXV, XXXVI and XXXVII, following the naming scheme of Sebé-Pedrós et al. (2014)). Myosin XXXVII is unique for Rhizaria, and marks the first known synapomorphy for the group (Figure 9). It is highly supported (100 % bs and 1.0 pp) and has a molecular signal distinct from other described myosins (Richards & Cavalier-Smith 2005, Sebé-Pedrós et al. 2014). In this rhizarian-unique class there has been an additional radiation within the chlorarachniophytes into three separate paralogs, all fully supported (100 % bs and 1.0 pp, Figure 9). Rhizarians have also gained a large repertoire of myosin IV, with six paralogs in Chlorarachniophyta and two in Foraminifera. All paralogs were well supported phylogenetically (bs > 90 % and pp >0.9) and differed from each other in functional domains (Figure 9). There was also a class unique to Chlorarachniophyta that resembled myosin IV by having a MYTH4 domain at the C-terminal, but with additional domains at the N-terminal usually not present in myosin IV (Richards & Cavalier-Smith 2005, Sebé-Pedrós et al. 2014). However both paralogs of Chlorarachniophyta in this class were phylogenetically distinct from myosin IV to warrant them to be given a new class (myosin XXXV). We also found myosin If in Chlorarachniophyta, which have been suggested to be present in the last common ancestor of all eukaryotes and recently lost in Rhizaria (Sebé-Pedrós et al. 2014). However the presence of two paralogs of myosin If in chlorarachniophytes show that it was present in the ancestor to Rhizaria, but that it might have been lost in Endomyxa and Retaria after they separated from the chlorarachniophytes. Finally there was a group unique to Chlorarachniophyta with two paralogs, named myosin XXXVI (Figure 9).

**Figure 9.**
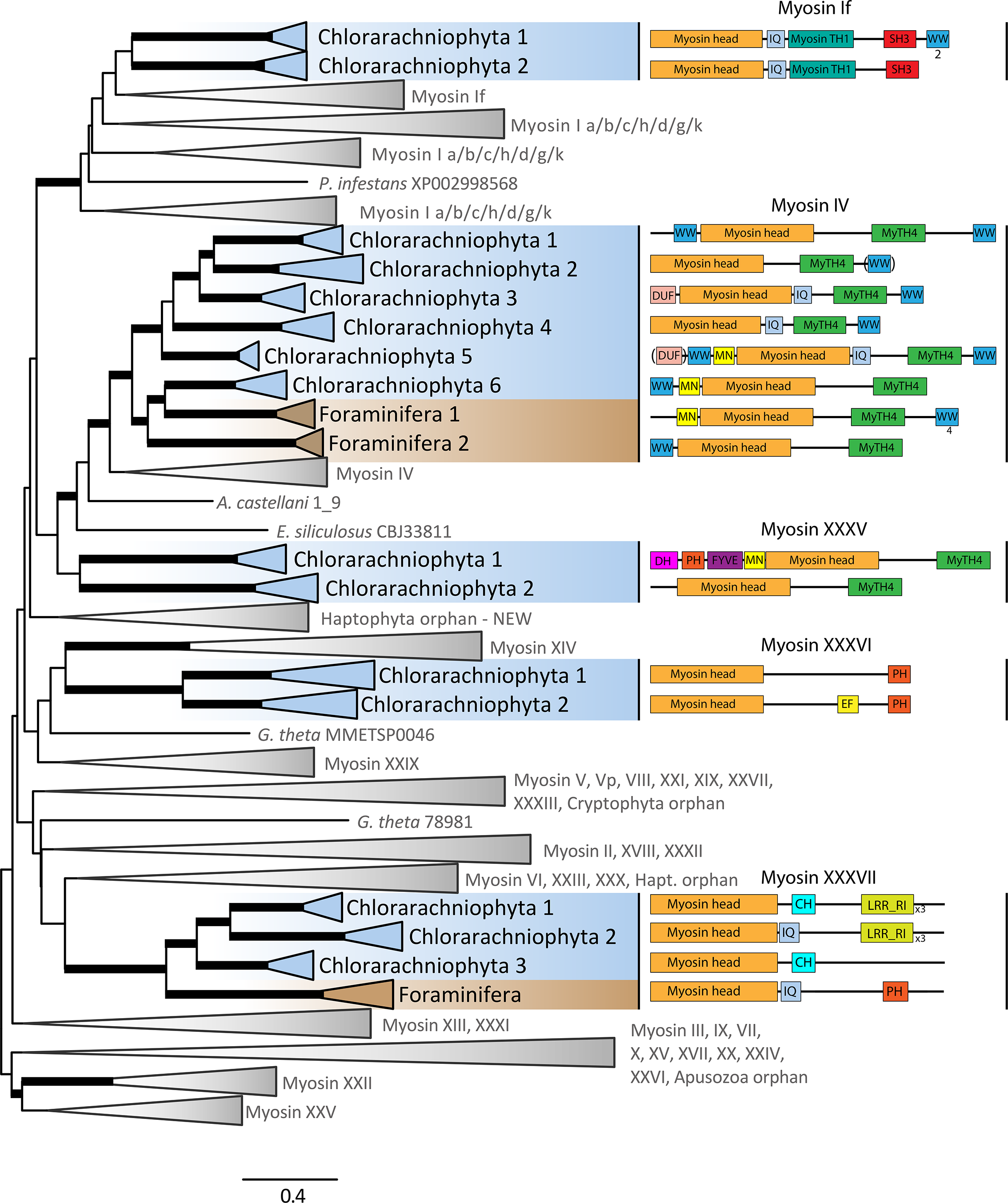
Myosin maximum likelihood phylogeny, (830 taxa, 754 AA). Groups coloured according to taxonomic affinity as in figure 2 (blue= Chlorarachniophyta, brown= Foraminifera). Branches collapsed according to myosin class affiliation and following the nomenclature of Sebé-Pedrós et al., (2014). The tree is midpoint-rooted and thick branches represents bootstrap > 75%, and Bayesian support > 0.8 pp. The scale bar equals the mean number of substitutions per site. The domain architectures for each class with representatives from Rhizaria are shown. IPR annotation of functional domains is listed Figure 9-figure supplement 1. A complete ML tree without collapsed branches can be found in Figure 9-figure supplement 2.

### α- and β-tubulin gene duplications in Retaria

We report 16 new α-tubulin and 19 new β-tubulin sequences from our two retarian transcriptomes: 4 α-tubulin, and 9 β-tubulin from *L. setosa*, 12 α-tubulin and β-tubulin from *S. zanclea*. Additionally, we identified 26 α-tubulin and 42 β-tubulin sequences from other rhizarian species in the MMETSP data (i.e. 12 Chlorarachniophyta and 14 Foraminifera α-tubulins; 10 Chlorarachniophyta and 32 Foraminifera β-tubulin). All these genes and other homolog sequences identified in GenBank were added to the Pfam seed alignment for α and β-tubulin (Finn et al. 2014). The phylogenetic tree of rhizarian α-tubulin revealed two different version of the gene: the canonical version of the α-tubulin gene (α1-tubulin; α1) and a novel group (α2-tubulin; α2) found only in Retaria (Figure 10). The split separating the two versions received maximal support (100% bs/ 1.0 pp). We identified α2-tubulin in the SCTs from both *L. setosa* and *S. zanclea*. Together with the available data from other Rhizaria, we confirm that this paralog is unique for Retaria. In Foraminifera there were several paralogs of α2, with most copies in *Reticulomyxa filosa* (25 copies). Foraminifera α2 was paraphyletic with a clade branching at the base of Retaria (100% bs/ 1.0 pp), before the Radiolaria and Taxopodida α2 clade. The bootstrap support was low (70 % bs), while the posterior probability was high (0.97 pp) for the branch that separated this group from the rest of the Foraminifera (Figure 10).

**Figure 10.**
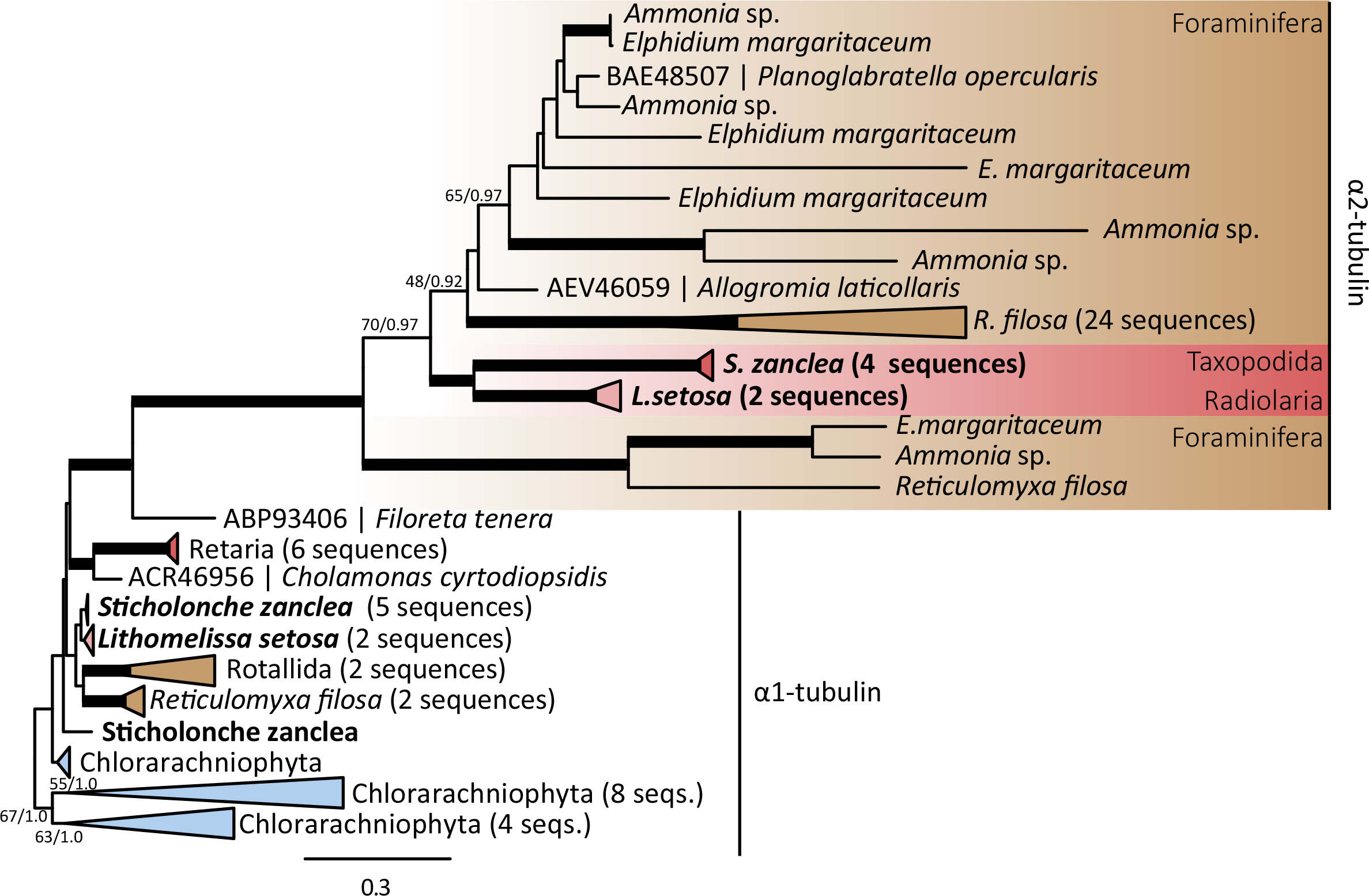
Phylogeny of rhizarian α-tubulin (75 taxa, 453 AA). Thick branches represents bootstrap > 75% and posterior probability > 0.9. Some branches are collapsed to save space. Support values for selected nodes discussed in the text are added for clarity. The colouring scheme is the same as in figure 2. (Brown = Foraminifera, red= Radiolaria, yellow = Endomyxa, and blue= Filosa). The scale bar equals the mean number of substitutions per site.

Similarly, the β-tubulin trees contained a clearly divergent clade (i.e. β2-tubulins) with several copies for each Retarian group (Figure 11). All β2-tubulin copies where grouped together with high support (100% bs/ 1.0 pp) in agreement with earlier studies (Hou et al. 2013). We also found that the β2 copies were present in Taxopodida as well as in Foraminifera and Radiolaria.

**Figure 11.**
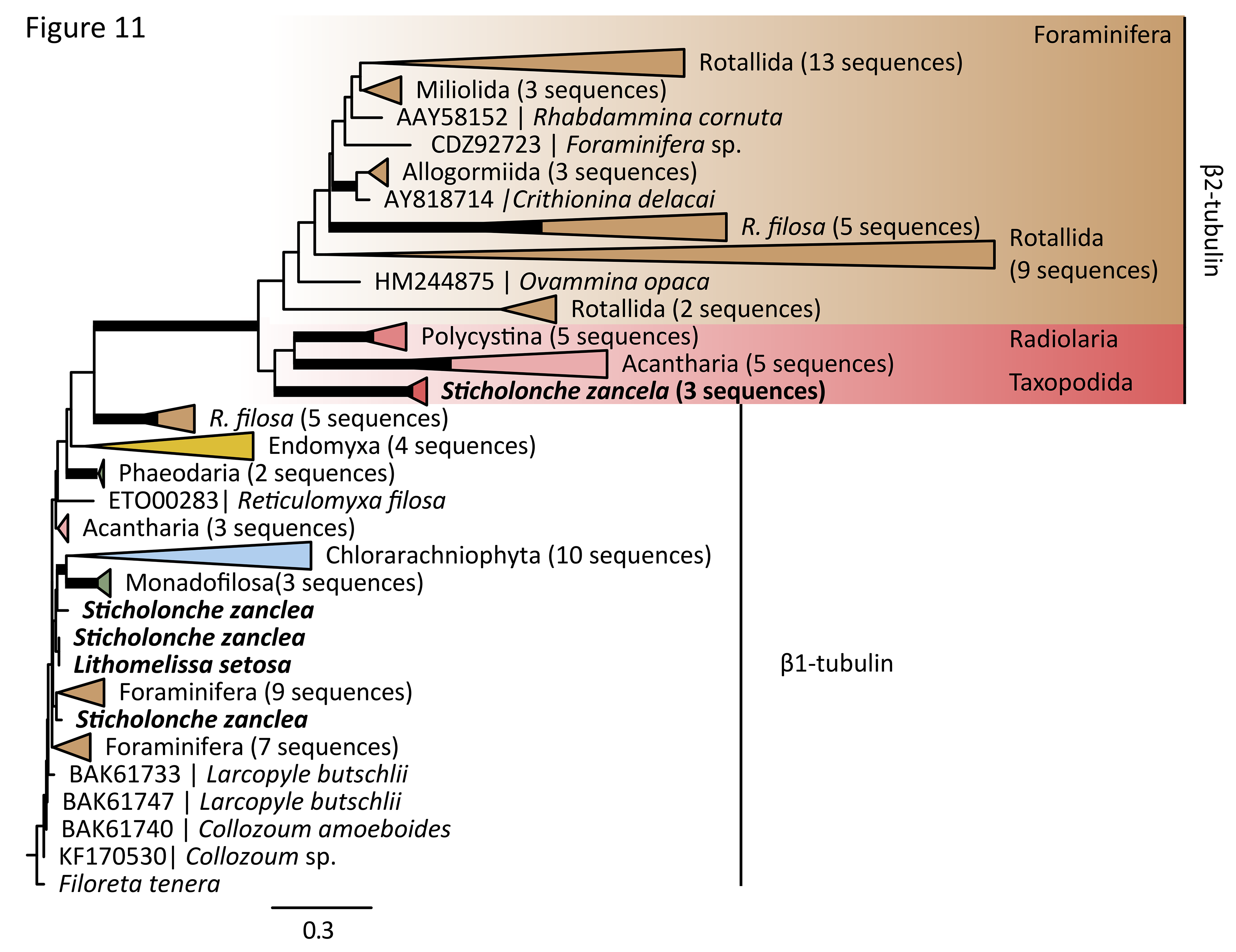
Phylogeny of rhizarian β-tubulin (104 taxa, 456 AA). Thick branches represents bootstrap > 75% and posterior probability > 0.9. Some branches are collapsed to save space. Support values for selected nodes discussed in the text are added for clarity. The colouring scheme is the same as in figure 2. (Brown = Foraminifera, red= Radiolaria, yellow = Endomyxa, and blue= Filosa). The scale bar equals the mean number of substitutions per site.

### Neofunctionalization of tubulin genes in Retaria

Comparative evolutionary analyses of the tubulin paralogs were done to identify patterns of functional change. This was performed by estimation of evolutionary rates and mapping site rates to tubulin structural models with Consurf (Ashkenazy et al. 2010, Celniker et al. 2013). Highly conserved amino acid residues were assumed to be functionally important and variable residues to be of less importance for function. We therefore compared separately α1 with α2, and β1 with β2; identifying sites conserved in one paralog and variable in the other. Such sites were believed to have undergone functional shifts and therefore considered important for cytoskeleton evolution. We also examined regions of the α- and β-tubulin structures known to be important for microtubule function and dynamics. Evolutionary changes in these areas are likely to affect the overall function of the microtubules.

Tracing evolutionary rates on the molecular structure of α- and β-tubulin (figure 12 and 13) revealed two patterns of functional change between the conventional and new tubulin genes: First, areas that are considered to be functionally important and conserved in α- and β-tubulin in general were conserved in α1 and β1 genes while being highly variable in α2 and β2, with differences being most prominent in α-tubulin. This pattern was observed for both inter-and intra-dimerization surfaces between monomers (i.e. longitudinal interactions important for protofilament assembly and disassembly), as well as lateral interactions between protofilaments. Both the T7-loop and H8 helix, important for longitudinal interactions, are extremely conserved in the original variant, while highly variable in the novel paralog. And similarly for the lateral contact points, helix H12, which forms a ridge on the outside of the microtubule and therefore affects binding and movement of motor proteins along the filament (Löwe et al. 2001) is much more variable in the novel α2-tubulin paralog than α1. Even the highly conserved residues in the M-loop of the original tubulin variants are highly variable for α2 and β2. These are residues directly interacting between neighboring protofilaments (Löwe et al. 2001). Taken together, all major contact areas both for lateral and longitudinal interactions are less conserved in the novel paralogs compared to the original, especially for α2.

**Figure 12.**
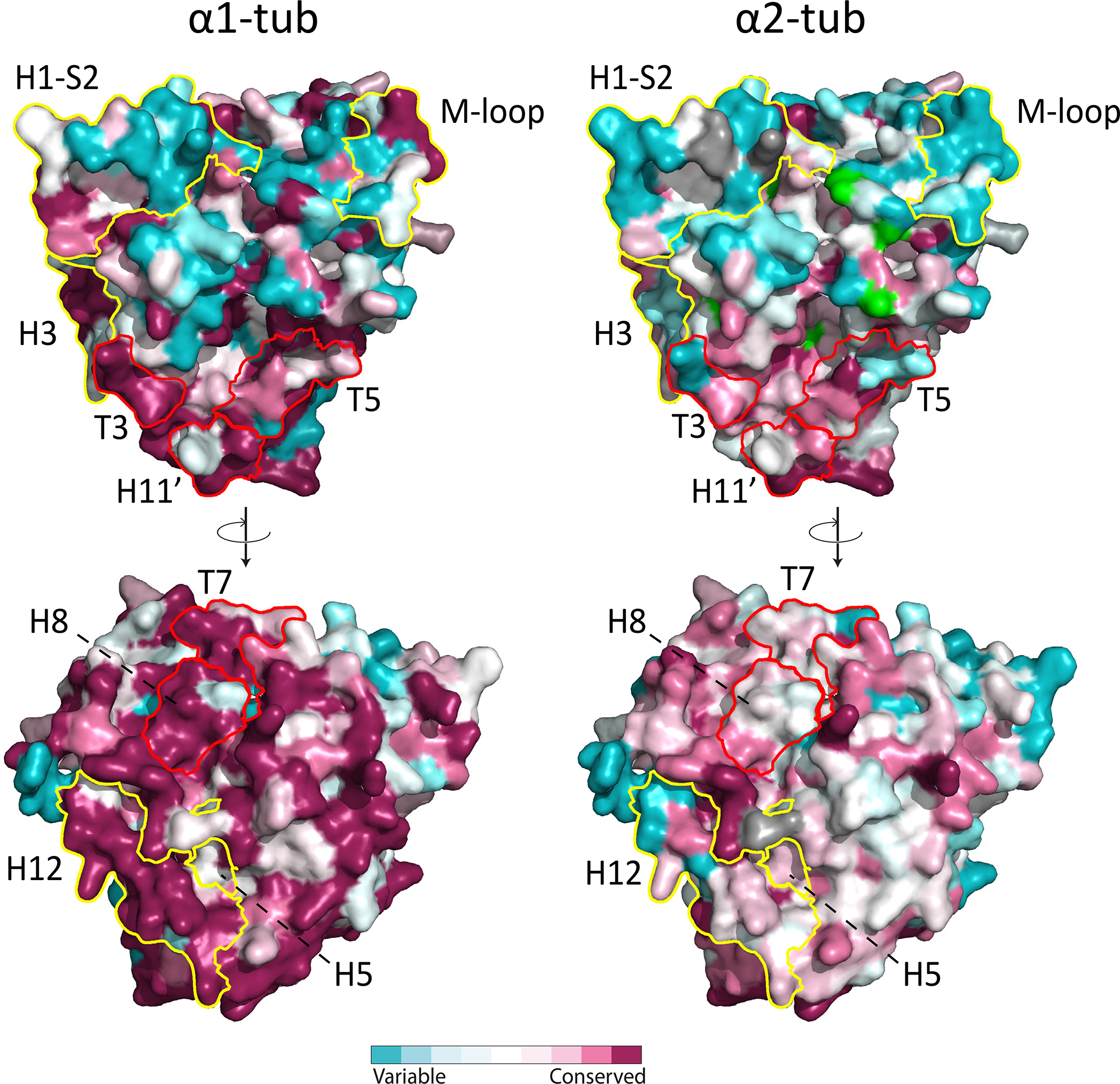
Molecular models of paralogs of α-tubulin (α1-and α2-tub) in Retaria using PDB accession 3du7 as template. Residues are coloured according to the evolutionary rates calculated by Consurf. Turquoise residues are highly variable and maroon means conserved residues.

**Figure 13.**
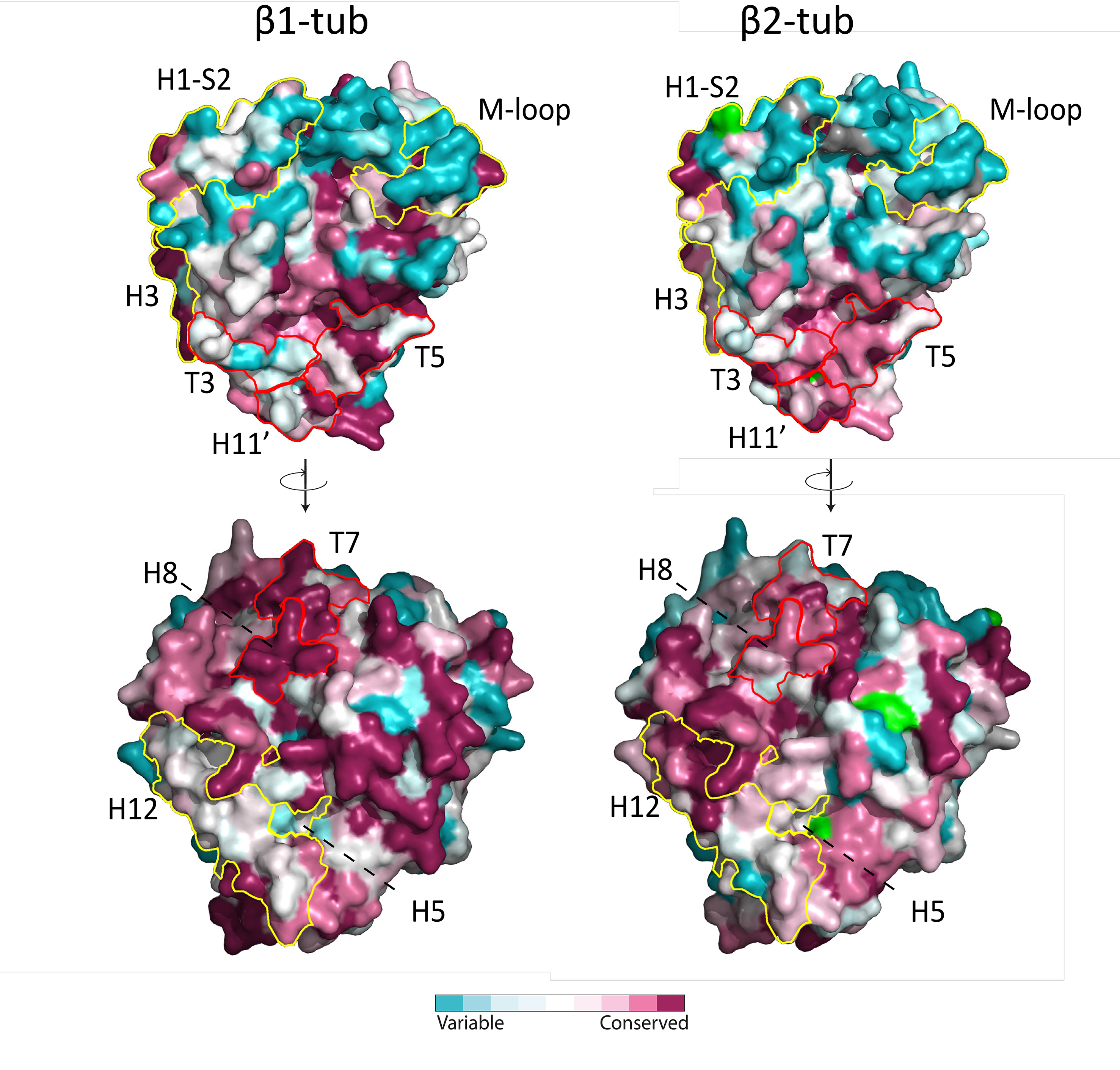
Molecular models of paralogs of β-tubulin (βi-and β2-tub) in Retaria using PDB accession 3du7 as template. Residues are coloured according to the evolutionary rates calculated by Consurf. Turquoise residues are highly variable and maroon means conserved residues.

Second, and in contrast to the pattern above, areas of the tubulin molecules considered to be functionally less important typically evolve faster than the contact surfaces. Several residues outside of the conventional longitudinal and lateral binding sites are highly conserved in both α2 and β2 while highly variable in the original α1 and β1 genes (Supplementary alignment). Many of these residues are exposed on the surface of the monomers and could represent new sites for other tubulin interactions or surfaces for motor protein attachment and movement.

Altogether, both the α2- and β2-tubulins have undergone dramatic evolutionary changes and are likely functionally distinct from their α1 and β1 counterparts (Figure 12 and 13).

## Discussion

The last common ancestor of Rhizaria was most likely a naked, heterotrophic flagellate, who relied extensively on its pseudopodia to explore the environment and to catch prey (Cavalier-Smith 2009). Its pseudopods were supported by actin and at least one group of myosins unique to Rhizaria (Figure 4 and 9, summarized in Figure 14). The Rhizarian cytoskeletons have since undergone evolutionary changes and their diversification follows a pattern where the major groups have their own favoured filament: the chlorarachniophytes have relied on actin to support their reticulose pseudopodia while the axopodia and reticulopodia in Retaria have been stiffened by microtubules composed of tubulin. Although some structural differences between lineages are known, little is established about the genetic basis of these phenotypes. Here we investigate the genetic evolution responsible for diversification of the cytoskeleton and present a hypothesis that relate these evolutionary changes to the varying morphology of Rhizaria groups. In order to fully understand the evolution of the cytoskeleton in Rhizaria we started by constructing a robust phylogenetic tree on which to map the evolutionary events.

**Figure 14.**
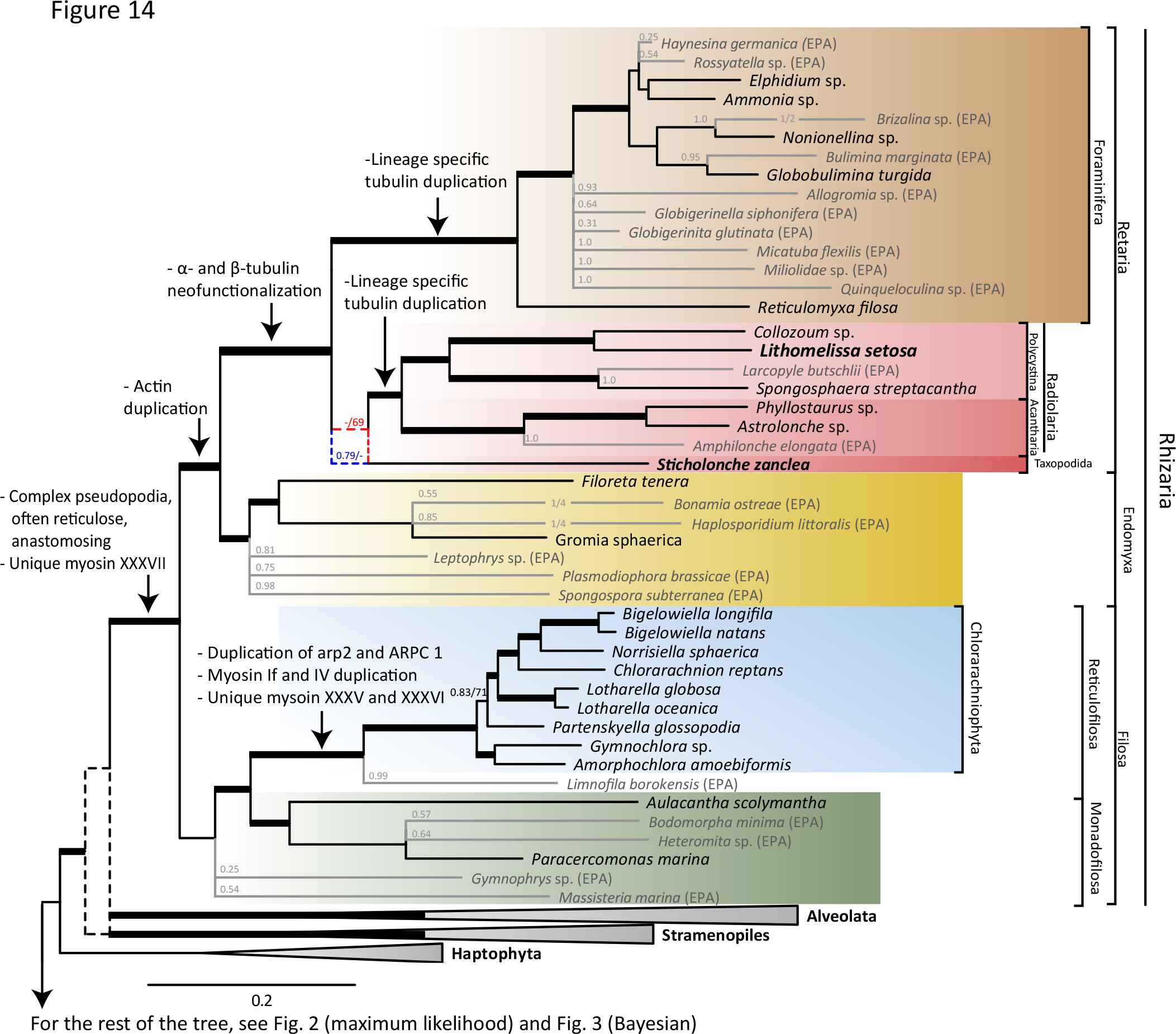
Phylogenetic tree of Rhizaria based on the full dataset, 255 genes, summarizing the major evolutionary events. Taxa with large portions of missing data are placed on the maximum likelihood reference tree with the Evolutionary Placement Algorithm (EPA; Berger et al., 2011). Taxa in bold are sequenced for this study. Arrows mark important evolutionary events, morphological changes and gene duplications. Thick branches are highly supported withe bootstrap support >90% and posterior probability > 0.9. Branches in grey are the most likely placement of taxa from EPA with numbers showing the expected likelihood weights for the placement. Branches that differ between maximum likelihood and Bayesian trees are marked with dashed lines. For *Sticholonche zanclea* the blue line represent the Bayesian CATGR placement with posterior probability, red dashed line represents the maximum likelihood LG placement with bootstrap support. For a further discussion of the placement of *Sticholonche zanclea* and the relationship between Alveolates, Stramenopiles and Rhizaria see the text. The scale bar equals the mean number of substitutions per site.

### Placing Rhizaria in the Tree of Life

Rhizaria has an evolutionary origin at the intersection between two very morphologically diverse and abundant lineages in the Tree of Life: the alveolates and stramenopiles. Together the three lineages form the supergroup SAR (Burki et al. 2007). Although there is little doubt that these three lineages are closely related to each other, the exact relationship between them remains debated. Three main hypotheses exists: either Stramenopiles and Rhizaria are monophyletic and sister to Alveolates (Burki et al. 2007, 2013, Katz & Grant 2014), alveolates and stramenopiles constitute a monophyletic group with Rhizaria as sister (Burki et al. 2008, 2010, 2012, 2016, Parfrey et al. 2010, Cavalier-Smith et al. 2015), or finally Rhizaria and Alveolates are monophyletic with Stramenopiles as sister group (Sierra et al. 2013, 2015, He et al. 2016).

Our Bayesian and ML inferences resulted in two different phylogenies. The Bayesian tree inferred with the CATGTR model grouped alveolates and stramenopiles, whereas the ML tree with the LG model clustered alveolates with Rhizaria. This inconsistency was evaluated by removing fast evolving sites, which have been suggested to contain misleading phylogenetic information (Philippe et al. 2005, Townsend 2007, Cummins & McInerney 2011, Townsend et al. 2012).

Removing such sites, caused no changes in the Bayesian phylogeny of the SAR groups, but the ML analyses converged towards the Bayesian tree by grouping alveolates and stramenopiles (Figure 4A and Suppl. table S4). The CATGTR model is a better representation of amino acid substitution patterns than the LG model, because it takes into account substitution pattern heterogeneity (Lartillot & Philippe 2004). In addition, the Bayesian inferences were less affected by site selection and were always reconstructing essentially identical phylogenies with the CATGTR model, altogether strongly supporting the grouping of alveolates and stramenopiles with exclusion of Rhizaria.

One of the remaining challenges about the SAR phylogeny is to identify the closest sister group of SAR in the global phylogeny of eukaryotes. Recently it has been proposed that haptophytes together with Centrohelida make up the sister clade to SAR (Burki et al. 2016). This is congruent to our Bayesian and ML trees where the haptophytes branch at the base of SAR, at least prior to the removal of fast evolving sites. However, removal of fast evolving sites typically groups haptophytes together with cryptophytes at the base of the Archaeplastida, as seen in other recent multi-gene phylogenies (supplementary table S4; Parfrey et al., 2010; Brown et al., 2012; Katz & Grant, 2014; Cavalier-Smith et al., 2015). This shifting position between SAR and the Archaeplastida may reflect that haptophytes diverged at the base of both groups and therefore is a key group for understanding the origin and evolution of this huge diversity of eukaryotes.

### Resolving Rhizarian Relationships

Within Rhizaria, it has been suspected for some time that Foraminifera and Radiolaria are closely related, and they have therefore been grouped together as Retaria (Cavalier-Smith 2002, Moreira et al. 2007, Krabberød et al. 2011, Ishitani et al. 2011, Sierra et al. 2013). In phylogenies based on ribosomal DNA, Foraminifera groups within radiolarians, although this placement has been contested based on the aberrant nature of both the small (18S) and large (28S) subunit of ribosomal genes in Foraminifera (Pawlowski & Burki 2009, Krabberød et al. 2011). Recent phylogenomic analyses place Foraminifera either within Radiolaria implying Radiolaria to be a paraphyletic group (Burki et al. 2013, Sierra et al. 2013, 2015) or as sister to Radiolaria (Cavalier-Smith et al. 2015, Burki et al. 2016). However, these analyses lack two crucial pieces in the puzzle; representatives from Nassellaria, one of the major polycystine radiolarian orders, and *S. zanclea*, the only species of Taxopodida. Including both in combined 18S and 28S rDNA phylogenies, divided the Radiolaria in two main groups, Polycystina and Spasmaria, where the latter contained Taxopodida, but the position of Foraminifera was unresolved (Krabberød et al. 2011). Here, we have generated transcriptome data and protein sequences from both the missing Radiolaria groups in our multi-gene analyses, *L. setosa* (Nassellaria) and *S. zanclea* (Taxopodida). In addition, we have reduced the impact of missing data in earlier phylogenomic analyses (Sierra et al. 2013, 2015, Cavalier-Smith et al. 2015, Burki et al. 2016) by adding genes to Foraminifera and a substantially larger sampling of other Rhizaria species.

Using these data, our analyses always cluster Radiolaria, Foraminifera and Taxopodida into Retaria. We find that Radiolaria (excluding Taxopodida) is monophyletic (congruent with Cavalier-Smith et al. 2015). Endomyxa and Retaria form a monophyletic group, revealing Cercozoa as paraphyletic. But in our multi-gene alignments, as in those of Sierra et al. (2013), data from two important endomyxean clades (i.e. Haplosporida and Vampyrellida) are absent. However, we included representatives from the two clades on the ML tree with the Evolutionary Placement Algorithm (Berger & Stamatakis 2011) and they fall inside the endomyxean clade, strengthening the monophyly of Retaria and Endomyxa (Figure 14).

### Taxopodida and Endomyxa revealed as sister lineages to Foraminifera and Radiolaria

Taxopodida have previously been placed within Radiolaria (Nikolaev et al. 2004, Krabberød et al. 2011), but has two different positions in our trees dependent on the analysis. The Bayesian CATGTR trees show Taxopodida as the sister to Radiolaria and Foraminifera, while ML LG place the species as sister to Radiolaria. We assessed the basis for this discrepancy by running Phylobayes with the substitution model used in the ML analyses (i.e. the LG model). The resulting Bayesian LG tree placed Taxopodida as sister to Radiolaria – congruent with the ML tree – clearly demonstrating the impact of the model on the phylogeny. It should also be noted that removing fast evolving sites in the ML LG analysis changed the tree correspondingly by placing Taxopodida at the base of Retaria (Figure 4B). While all the Bayesian inferences were highly congruent, the ML topologies were less stable and converged towards the Bayesian tree with removal of fast evolving sites. The stability of the Bayesian results may be due to the use of the CATGTR model which more realistically estimates the evolutionary substitution patterns in amino acids by taking into account across site heterogeneities in the amino acid substitution process (Lartillot & Philippe 2004, Lartillot et al. 2013) and therefore preferable over the LG model.

All evidence taken into account, Taxopodida most likely diverged early in the radiation of Retaria and before the separation of Radiolaria and Foraminifera. This has consequences for the interpretation of the cytoskeleton and morphological evolution of Retaria. Taxopodida and Acantharia were grouped together as Spasmaria based on the existence of contractile myonemes in both groups (Cavalier-Smith 1993), a grouping also supported in 18S and 28S rDNA phylogenies (Krabberød et al. 2011). Myonemes give taxopodidans the ability to swim using their pseudopodia like oars while giving acantharians the ability to regulate their buoyancy by altering their cell volume (Cachon et al. 1977, Febvre 1981). However, if Taxopodida is sister to both Radiolaria and Foraminifera it implies that contractile myonemes and flexible pseudopodia, were an ancestral trait of Retaria, and have later been lost or modified in Radiolaria and Foraminifera.

Endomyxa was originally defined as a clade within Cercozoa (Cavalier-Smith 2002). In our trees, however, Endomyxa was consistently excluded from the filose Cercozoa in both ML and Bayesian inferences, and placed as sister to Retaria. Our trees show both Endomyxa and Taxopodida as sister lineages to Foraminifera and Radiolaria. This means that Rhizaria is split into three lineages: Filosa, Endomyxa and Retaria. Taxopodida, Foraminifera and Radiolaria constitute Retaria. This new branching order of rhizarian lineages forms the framework we here use to map changes of the cytoskeleton-related gene families and establish the order of macroevolutionary changes in Rhizaria.

### Expansion of actin, myosin and subfunctionalization of Arp2/3 in Chlorarachniophyta

The chlorarachniophytes can form extensive networks of reticulose actin-based pseudopodia that they rely on for foraging and movement (Margulis 1990). The evolution of these extensive pseudopodial networks seems to have been made possible by gene duplications of proteins controlling actin network dynamics as well as several duplications of the actin gene, and of myosin specific to chlorarachniophytes. The interaction between actin, the Arp2/3 complex, and myosin is important for pseudopod formation and branching. Branching points between two actin filaments are formed as the Arp2/3 complex recruits actin filaments into networks (Volkmann et al. 2001, Goley & Welch 2006, Pollard 2007, Mattila & Lappalainen 2008, Xu et al. 2011). Here we present evidence for a duplication ancestral to chlorarachniophytes for two of the proteins in the complex: Arp2 and ARPC1. Both proteins are involved in the initial binding of nucleation promoting factors (NPFs) that are essential for the formation of protrusions that eventually leads to pseudopodia at the leading edge of motile cells (Boczkowska et al. 2008, 2014, Xu et al. 2011, Ura et al. 2012, Kast et al. 2015). Although the exact nature and conformation of the Arp2/3 complex are still under investigation, it seems clear that actin NPFs bind first to Arp2 and ARPC1, then extend the daughter filament by adding an actin subunit at the barbed end of Arp2 and Arp3 (Boczkowska et al. 2008, 2014). This in turn creates attachment points for daughter actin filaments to bind to the existing mother filament (Rouiller et al. 2008). In chlorarachniophytes the Arp2 and ARPC1 paralogs have undergone divergent substitution patterns. The differences between the two Arp2 paralogs as well as the two ARPC1 paralogs are mainly found on the surface areas of the Arp2/3 complex where the actin recruiting proteins, NTPs and ultimately the newly formed actin filaments attach. Sites that are conserved and shared between both paralogs (marked green in Figure 8) are most likely important for the original function of the complex, while the sites that are conserved in one of the paralogs but not the other points to functional differentiation and innovation. In addition myosin duplications have occurred ancestrally to Rhizaria before several independent events in chlorarachniophytes and Foraminifera.

Over evolutionary time scales these genetic innovations have likely formed the molecular basis of cellular and morphological differentiation in chlorarachniophytes: In turn, this has given them a larger repertoire of Arp2 and ARPC1 and an increased potential to recruit actin filaments to facilitate a reticulate cell and a gliding lifestyle.

### Unique duplication and neofunctionalization of α- and β-tubulin in Retaria

Similar to Chlorarachniophytes many species in Retaria and Endomyxa can form highly branched pseudopodial networks (Anderson 1976a, b, Lee & Anderson 1991, Suzuki & Aita 2011). This is also reflected in the expansion of actin genes: Retaria has two distinct subfamilies of actin genes, one grouping with actin homologs from Endomyxa. In Endomyxa the actin diversity is extensive with three possible paralogs (Figure 4). Unlike Chlorarachniophyta however, Retaria have additional pseudopods supported by microtubules called axopodia (Anderson 1983, Travis & Bowser 1986, Lee & Anderson 1991, Suzuki & Aita 2011). The axopodia in Radiolaria are often contractile and withdraw upon contact; rapid movement can cause prey to be drawn towards the cytoplasm of the cell where digestion occurs (Sugiyama et al. 2008). Similarly Foraminifera have stiffened pseudopods called reticulopodia. These microtubule mediated pseudopods can extend and retract at a speed two orders of magnitude faster than in animal cells (Travis & Allen 1981, Bowser 2002). The extraordinary speed at which the microtubules can nucleate in Foraminifera has been linked to a duplication and neo-functionalization of β-tubulin (Habura et al. 2005, Hou et al. 2013). The discovery of the aberrant β2-tubulin was a paradox, because the corresponding α-tubulin paralog of the heterodimer was absent in all Retaria (Hou et al. 2013). The question is how an aberrant β-tubulin can function without a correspondingly deviant α-tubulin. Here we solve this paradox by presenting α2-tubulin in the single cell transcriptomes of *Sticholonche zanclea* and *Lithomelissa setosa*, which enabled identification of homologs from other Retarian species. We also add new α-tubulin data from both *S. zanclea* and *L. setosa*, confirming gene expansion in all major Radiolaria lineages, and the origin of new paralogs in the common ancestor to Retaria. Interestingly, none of the α2-tubulin and β2-tubulin paralogs could be identified in available Endomyxa data, suggesting that these gene duplications are synapomorphic for Retaria, with an origin after Retaria and Endomyxa diverged (Figure 14).

Both the α2- and β2-tubulin genes form distinct phylogenetic clades and diverged from the conventional α1-and β1-tubulin genes through changes in functionally important residues. Subsequent to the initial duplication, these paralogs have undergone repeated duplications to form subgroups of each gene family. These duplications and increased evolutionary rate have developed both α2- and β2-tubulin as more divergent genes than the conventional α1-and β1-tubulins.

Modelling of evolutionary rates on the tubulin structure shows global changes of the molecule along two different paths: Firstly, a large number of conserved and functionally important residues in α1 and β1 have become more variable, and probably therefore less functionally important in α2 and β2. This pattern is particularly clear at the interface between the α and β heterodimers (which is the basic unit of protofilaments), and in the lateral surfaces between protofilaments that create microtubule (i.e. the M-loop, the T3-loop and 8H helix etc.). Secondly, many variable sites localized outside of the classical contact surfaces in the conventional α1 and β1, have become conserved in α2 and β2 and have probably gained new functional roles. In addition, tracing the evolutionary rates of all tubulin paralogs show higher evolutionary rate at the interface between the α and β heterodimers (which are the basic units of protofilaments), and in the lateral surfaces between protofilaments that create microtubules (i.e. the M-loop, the T3-loop and 8H helix etc.). The overall pattern is that the new α2-tubulin paralog presented here evolved with a similar mode to that of the β2-tubulin gene (Habura et al. 2005, Hou et al. 2013).

Retaria is unique among eukaryotes in having such divergent tubulin genes. It is not clear how Retaria combines the four tubulin variants α1, α2, β1 and β2 into heterodimers, but it certainly enables modularity. We hypothesize that Retaria can assemble four types (type 1-4) of heterodimers; i.e. *type1:* α1+β1, *type2:* α1+β2, *type3:* α2+ β1, *type4:* α2+ β2. These four heterodimers can function as modules and can be combined to develop protofilaments with different properties. The different affinities between the α and β tubulins will certainly affect assembly and disassembly of microtubules, and may be used to adjust flexibility, strength and conformation of the axopodia or reticulopodia (Löwe et al. 2001). Retaria is known to develop elaborate pseudopodia stiffened by bundles of microtubules. The axopodial microtubules in Taxopodida and Acantharia attach laterally to other microtubules and form multiple hexagonal rings (Cachon et al. 1973). The microtubules of polycystine radiolarians axopodia on the other hand form a branching pattern (Cachon & Cachon 1971, Grain 1986). In some Foraminifera (e.g. *Astrammina*) the microtubules coil tightly around one around another increasing the tensile strength of the pseudopodia used to capture prey (Lee & Anderson 1991). Having several types of α and β heterodimers allows a large repertoire of architectural structures to be drawn upon when forming microtubules. In addition, we observe that many of the sites that have undergone evolutionary change are located on the surface of the heterodimer. This can be linked to binding sites for microtubule associated proteins (MAPs) as well as motor proteins further expanding the range and flexibility of cytoskeletal structures (Brouhard & Rice 2014).

### Single cell transcriptomics for macroevolutionary studies of unculturable protists

Here we have applied single cell transcriptomics as a new approach for phylogenomic reconstruction of SAR, and the genetic basis of cytoskeleton and morphological evolution in Rhizaria. Presently single cell transcriptomics has been applied to animal model organisms, such as cell differentiation studies in mouse (Liang et al. 2014, Liu et al. 2014). To date, the only application on protists has been on single cells grown from culture, confirming that the method gives comparable results to that of sequencing many thousand cells (Kolisko et al. 2014). Although single cell transcriptomic studies from cultured cells confirm the approach, they do not address how efficient the method works on free-living protists, with highly divergent cell types, from natural samples.

One of the main challenges of applying single cell transcriptomics to protists is the optimization of cell lysis. This is emphasized when studying species with rigid skeletons and tough cell walls. Here we modified lysis procedures for single cell transcriptomics (Picelli et al. 2014). Radiolaria species have a though cellular wall that protects the endoplasm; successful lysis demonstrates how the method can be applied to less hardy unicellular species. The number of predicted genes from our single cell transcriptomes are comparable to that generated from colonies, or pooling of hundreds of cells from other Radiolarian species (Burki et al. 2010, Balzano et al. 2015). Subsampling of sequence reads showed sufficient sequencing depth, suggesting that an incomplete transcriptome was likely due to stochastic loss of mRNA. Despite these challenges we have shown that transcriptomes of sufficient quality for phylogenomic and molecular evolutionary analyses can be generated from single cells isolated from natural samples. This protocol can undoubtedly be applied to other uncultivable protists, adding resolution to the relationships between eukaryotes, in addition to revealing the evolution of morphologically related genes.

### Morphological diversification by lineage specific innovation of cytoskeleton genes in Rhizaria

Data generated from these transcriptomes demonstrate that genetic innovation through multiple gene duplication and neo-functionalization processes, rather than co-option of deep gene homologs, have taken place in cytoskeletal genes of Chlorarachniophyta and Retaria. Differential expansion of genes in chlorarachniophytes and Retaria show that underlying genetic changes to cytoskeletal evolution have taken different routes in morphologically distinct groups; the overall pattern of the data reveals extensive gene duplications of actin-related proteins in chlorarachniophytes and of α- and β-tubulins in Retaria, with group specific expansions of myosin in both groups (Figure 14). The hypothesized connection between the evolutionary changes to cytoskeletal genes and the cellular morphology of the cells suggest that genetic innovations occurred in the ancestor of the respective groups, subsequently forming the basis for morphological and species diversification. While the actin-related proteins, and the myosin motor proteins that use them have driven changes in chlorarachniophytes; tubulin has directed central components of Retaria evolution. Subsequent to the initial innovation, additional expansions of functional genes crucial to cytoskeletal formation have impacted on the morphological diversification of Chlorarachniophyta and Retaria. Our analyses elucidate relationships between genotype and phenotype of these organisms, linking gene evolution to evolution of cell morphology. Better understanding of macroevolution in these organisms will require functional studies of what types of actin branching the new Arps can form in chlorarachniophytes and how Retaria combine the two sets of α and β tubulin proteins in their protofilaments. Such studies should be complemented with more data from other gene families known to be involved in cytoskeleton development, regulation, and transportation, such as MAPs, GTPases, dynein and kinesin (Hammer & Wu 2002, Kollmar et al. 2012, Rojas et al. 2012, Brouhard & Rice 2014),.

Using the transcriptome data, we addressed uncertainties in the phylogeny of SAR by phylogenomic analyses of supermatrices and by identification of synapomorphic gene duplications. The phylogenomic analyses strongly support Rhizaria as sister to Stramenopiles and Alveolates, and reveal Endomyxa and Taxopodida as two sister lineages to Foraminifera and Radiolaria, with the latter two being divided into two distinct clades. Foraminifera is not placed within Radiolaria as earlier reported (Krabberød et al. 2011, Sierra et al. 2013, 2015). In addition, we identified independent synapomorphic gene duplications characters on several taxonomic levels in Rhizaria, including a myosin family unique to Rhizaria. The actin-2 paralog of Foraminifera, Radiolaria and Taxopodida is shared with Endomyxa, arguing for the grouping of Endomyxa with Retaria, instead of being placed within Cercozoa. However, Retaria is divided from Endomyxa by the Retaria-specific α2 and β2 tubulin synapomorphies. The duplication of genes in the Arp2/3 complex is a synapomorphy for Chlorarachniophyta. Altogether the phylogenomic trees and synapomorphic gene-duplications shown here, form a new framework for future revisions of the classification of SAR and Rhizaria. In total, we demonstrate single cell transcriptomics as a promising approach for inclusion of a larger diversity of uncultured protists in macroevolutionary studies and phylogenomic inferences.

## Methods

### Sampling and transcriptome amplification

Plankton samples were collected from the inner part of the Oslo fjord (May 2014) using a net haul with mesh size of 60 μm. The seawater samples were stored overnight in an incubator holding the same temperature as the fjord to let living cells recover and self-clean. Radiolarian cells were manually extracted from the plankton samples by capillary isolation with Pasteur pipettes and an inverted microscope. Cells were individually photographed and then thoroughly washed in sterile PBS to remove possible surface contamination (Figure 1). Immediately following isolation, cells were placed in Nucleospin RNA XS lysis buffer (Macherey-Nagel) and processed further. Total RNA was isolated from the free-living Radiolarian cells using Nucleospin RNA XS (Macherey-Nagel) following standard protocol, with on-column DNase treatment and eluting with 5μl elution buffer. Hybridization of oligo(dT) primer, reverse transcription, template switching and PCR amplification of cDNA were performed by modification of a protocol outlined in (Picelli et al. 2014) called Smart-seq2; we used 7 μl of mRNA mix (5 μl isolated RNA, 1 μl oligo(dT) primer and 1 μl 10 mM dNTPs) which was added to 9 μl of reverse transcriptase (RT) mix). All 16 μl (mRNA+RT mix) was used for PCR amplification (adjusting concentrations accordingly) employing 20 cycles. The quality and integrity of the resulting cDNA were confirmed using a Bioanalyzer (Agilent) with a high-sensitivity DNA chip, in addition to visualization on a 1% TAE gel. cDNA concentration was confirmed using a Qubit fluorometer (Life Technologies) and the dsDNA HS assay kit.

### Sequencing and assembly

Library preparation and sequencing of the cDNA with Illumina MiSeq were performed at the Natural History Museum in London. The sample was prepared using the Illumina TruSeq Nano DNA LT Library Preparation Kit (FC-121-4001). The standard Illumina protocol was followed with fragmentation on a Covaris M220 Focused-ultrasonicator. The finished library was quality checked using an Agilent Tapestation to check the size of the library fragments, and a qPCR in a Corbett RotorGene instrument to quantify the library. This was repeated for two MiSeq 600 cycle runs, 2*300 cycle paired end sequencing. The MiSeq platform was chosen over HiSeq since the longer reads would provide an easier assembly when dealing with a possible metatranscriptomic library. The raw reads (19,894,654 for *S. zanclea* and 11,590,658 for *L. setosa*) were quality filtered and pairwise assembled with PEAR (Zhang et al. 2013) using default parameters. The reads where further cleaned with Trimmomatic (Bolger et al. 2014) and then *de novo* assembled into contigs with Trinity (Haas et al. 2013) using default settings. TransDecoder in the Trinity package was used to predict genes from the assembled cDNA (Haas et al. 2013).

To check if all transcripts in the library had been sequenced, the raw reads were randomly split up in 10 different datasets representing 10%, 20%, up to 90% of the original raw reads. The subsampled datasets were assembled and new gene predictions were independently performed using PEAR, Trimmomatic and Trinity as for the full dataset. Accumulation curves obtained by plotting the predicted gene number against increasing partition size show that the slope of the curves decrease with increasing partition size and more or less flattens when it reaches 100% of the total dataset for both libraries (Figure 1 – figure supplement 1). We therefore assume that acceptable sequencing depth for each library has been achieved, and that a further sequencing effort would not have increased the number of predicted genes significantly.

### Alignment construction, paralog identification, and phylogenetic inference

**The BIR pipeline:** We used the BIR pipeline (www.bioportal.no; Kumar et al., 2015) to extract genes and prepare single gene alignments to be used in multi-gene phylogenetic analyses. As seed alignments for the BIR pipeline we used 258 genes previously published in multi-gene phylogenies (Burki et al. 2012). As a query database we used the generated transcripts from our single cell transcriptomes (6898 in total), all proteins in GenBank with Rhizaria as TaxID (44278 sequences at the time of retrieval, October 2014), all 16 transcriptomes assigned to Rhizaria from the Marine Microbial Eukaryote Transcriptome Sequencing Project (MMETSP; Keeling et al., 2014. See table 1), as well as all Rhizarian sequences from Sierra et al., (2013). In addition seven reference genomes are included in the BIR pipeline (*Arabidopsis thaliana, Bigelowiella natans, Dictyostelium discoideum, Guillardia theta, Homo sapiens, Monosiga brevicollis, Naegleria gruberi, Paramecium tetraurelia, Saccharomyces cerevisiae and Thalassiosira pseudonana* (Kumar et al. 2015). In short the BIR pipeline will screen the query sequences against the database consisting of one or more seed alignments, using BLAST, and assign the sequences that match the criteria set by the user to the corresponding alignment (for details see Kumar et al., (2015)).

**Single gene analyses**: Maximum Likelihood (ML) trees for all single genes were constructed with RAxML v 8.0.2, with the program calculating the best fitting model for each gene (the option -m PROTGAMMAAUTO), and with the automatic bootstrapping criteria MRE (option -l autoMRE) (Pattengale et al. 2010, Stamatakis 2014). The Tree Certainty index (Salichos et al. 2014) was calculated for each tree separately, and all trees were run through a custom made R script to see whether the following clades were monophyletic or not: Opisthokonta, Fungi, Alveolata, Stramenopiles, Haptophyta, Rhizaria, Viridiplantae, Excavata, Fungi and Rhodophyta. This allowed us to screen for genes containing artefacts and dubious sequences such as sequences that had been assigned to the wrong species, sequences that originated from contamination and possible paralogs. Three genes (β-tubulin, actin and rac1) were found to have paralogs and deemed not suitable for multi-gene phylogenies. We therefore proceeded with 255 genes for the multi-gene analysis.

**Supermatrix construction:** After screening we were left with 255 genes that were concatenated using ScaFos (Roure et al. 2007). We also merged close species into composite sequences when they covered different parts of the supermatrix (see table S2). The final matrix had a length of 54,898 amino acids with 124 taxa.

**Removal of jumping and long branched taxa:** *Mikrocytos mackini* was not included in the analysis due to an extremely long branch (Burki et al. 2013) and RogueNaRok, using default parameters (Aberer et al. 2013) was used to identify jumping taxa, which also were excluded from further analysis (Supplementary table S2)

**Reduced dataset:** We also constructed a concatenated dataset consisting of 146 representative genes for easier and faster analysis. The selection of genes were made to meet several criteria: we excluded genes that had less than 45 taxa (50% of the inferred taxa), a low relative Tree Certainty index (Salichos et al. 2014), or that failed to group at least two of the major clades mentioned above.

**Missing data:** To assess the impact of missing data we excluded taxa with low coverage in increments from the two concatenated dataset. First we sat the lowest allowed percentage of missing data for a taxon to be 10% of the total characters (i.e. if a taxon had more than 90% data missing it was excluded), the next cut-off at 20%, and finally at 30%. The number of characters in the matrix was held constant. The Tree Certainty index (Salichos et al. 2014) was calculated for each increment see supplementary table S2 and S3 for details. The relative Tree Certainty index increased markedly when the threshold was set at 90%, but did not increase significantly after that, in fact there seem to be a decrease in the relative TC value as the number of taxa drops (Supplementary table S1).

**Influence of taxa with low coverage, or uncertain position**: we also remove taxa and clades from Rhizaria that had a consistently low bootstrap value (< 75%) or low posterior probability (< 75 pp), but that had not been flagged by RogueNaRok to see if they affected the topology of the phylogenetic inference. *Spongosphaera streptacantha* and *Sticholonche zanclea* were removed one by one and together from both the full and the reduced dataset. Foraminifera were also removed in analyses (see table S3).

**Removal of fast evolving sites:** TIGER (Cummins & McInerney 2011) was used with default settings to produce categories of fast evolving sites, 10 categories in total for each dataset. Categories of fast evolving sites were removed in increments, starting with the category with the fastest evolving sites, subsequently removing the category with the second fastest evolving sites etc. Up to 4 categories where removed from all datasets before phylogenetic analyses.

**Phylogenetic analyses:** Phylogenetic trees were inferred for all concatenated datasets, with RAxML choosing the best fitting model, and with the automatic bootstrapping criteria as previously described. The preferred model was always LG+r (see supplementary table S3). Due to the heavy demand on computational resources from Bayesian inference only six of the alignments were included for analysis with the CATGTR model in Phylobayes MPI version 1.5a (Lartillot et al. 2013), as well as 1 dataset with the LG model. For these we ran 2 chains in parallel for at least 15.0 iteration only stopping when the maxdiff was >0.3 (see supplementary table S3).

**Evolutionary Placement Algorithm:** In order to place rhizarian species that had been excluded when the cut-off threshold for missing data had been raised on the phylogenetic tree we used the Evolutionary Placement Algorithm (EPA) included in RAxML 8.0.26 (Stamatakis et al. 2010, Berger et al. 2011, Stamatakis 2014). As reference tree we used the 255-gene maximum likelihood tree with a 10% missing data cut off.

### Genes related to cytoskeleton formation and motor proteins

The assembled transcriptomes from the single cells were annotated with InterProscan 5 (Jones et al. 2014) as implemented in Geneious 8 (Kearse et al. 2012). The annotations were screened for genes commonly involved in the formation and development of the cytoskeleton, as well as the most common motor proteins using the cytoskeleton. In particular we looked for α- and β-tubulin, myosin, actin, the actin regulating Arp2/3-complex consisting of seven actin-related proteins (arp2, arp3, ARPC1, ARPC2, ARPC3, ARPC4 and ARPC5). Reference alignments and sequences were downloaded from PFAM (http://pfam.xfam.org/), as well as relevant other recently published alignments (Hou et al. 2013, Sebé-Pedrós et al. 2014, Cavalier-Smith et al. 2015) and used in BIR as seed alignments with the same query database as before. In addition representatives for all the genes where blasted against 6 additional non-rhizarian transcriptomes from MMETSP (MMETSP0039 *Eutreptiella gymnastica*, MMETSP0046 *Guillardia theta*, MMETSP0308 *Gloeochaete wittrockiana*, MMETSP0380 *Alexandrium tamarense*, MMETSP0902 *Thalassiosira Antarctica*, and MMETSP1150 *Emiliania huxleyi)*, as well as against the non-redundant protein database in GenBank. For each gene ML trees were constructed with RAxML as before, and manually curated for any confounding artefacts. Redundant and short sequences were manually removed in Geneious 8 (Kearse et al. 2012) before another round of ML analysis with RAxML and a Bayesian analysis with the CATGTR model implemented in Phylobayes MPI version 1.5a (Lartillot et al. 2013). Comparative evolutionary analyses of tubulin and the duplicated genes in the Arp2/3 complex were performed by examining the evolutionary rates of the paralogs separately and then mapping the genes to structural models using Consurf (Ashkenazy et al. 2010, Celniker et al. 2013). InterPro annotations of functional domains of myosin was performed with InterProscan 5 (Jones et al. 2014, Mitchell et al. 2015).

All sequences from *S. zanclea* and *L. setosa* used in this study have been deposited in GenBank with accession numbers (xxxx-xxxx). All data, alignments, and trees can be downloaded from www.bioportal.no.

## Acknowledgment

We would like to thank Bente Edvardsen, UiO, for providing the plankton haul from which the sampled cells were isolated, the Sequencing centre at Natural History Museum in London for performing the Illumina library preparations and sequencing. We would also like to thank Simon Picelli for answering questions related to the Smart-seq2 method, Fabien Burki for providing single gene alignments and The Gordon and Betty Moore Foundation for making all those wonderful protists transcriptomes available for the public. All analyses were run either on the Abel supercomputer at The High Performance Computing cluster at University Of Oslo, or on Lifeportal (www.lifeportal.uio.no). For more information on BIR see www.bioportal.no. This project was funded by University of Oslo. This work was supported by grants from University of Oslo and from Research Council of Norway to Shalchian-Tabrizi (NFR216475).

## SUPPLEMENTAL MATERIAL

**Figure 1-figure supplement 1:**
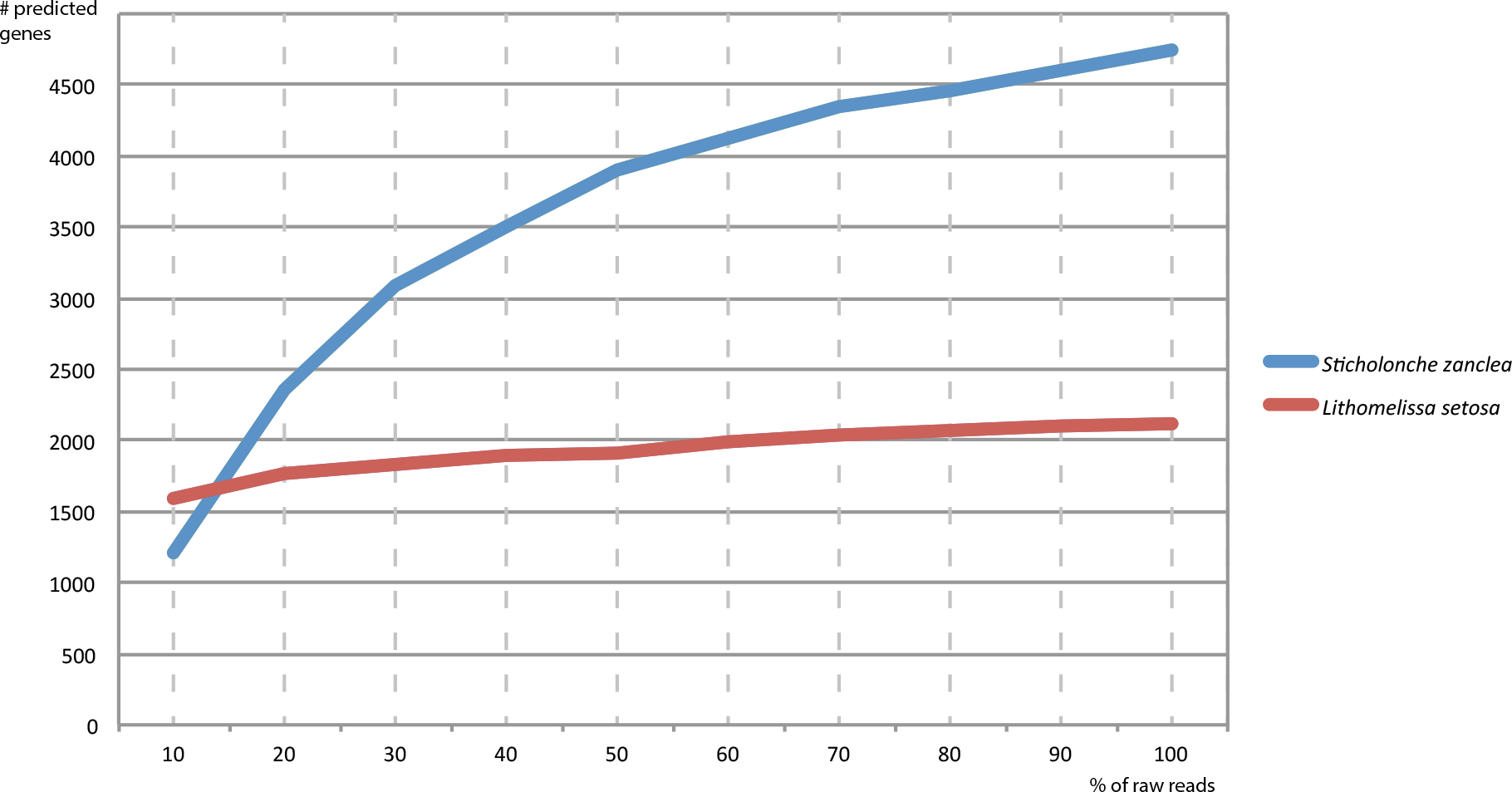
Gene accumulation curve. The number of predicted genes is plotted against subsamples of the original dataset. Each subsample is independently assembled before gene prediction.

**Figure 2-figure supplement 1:**
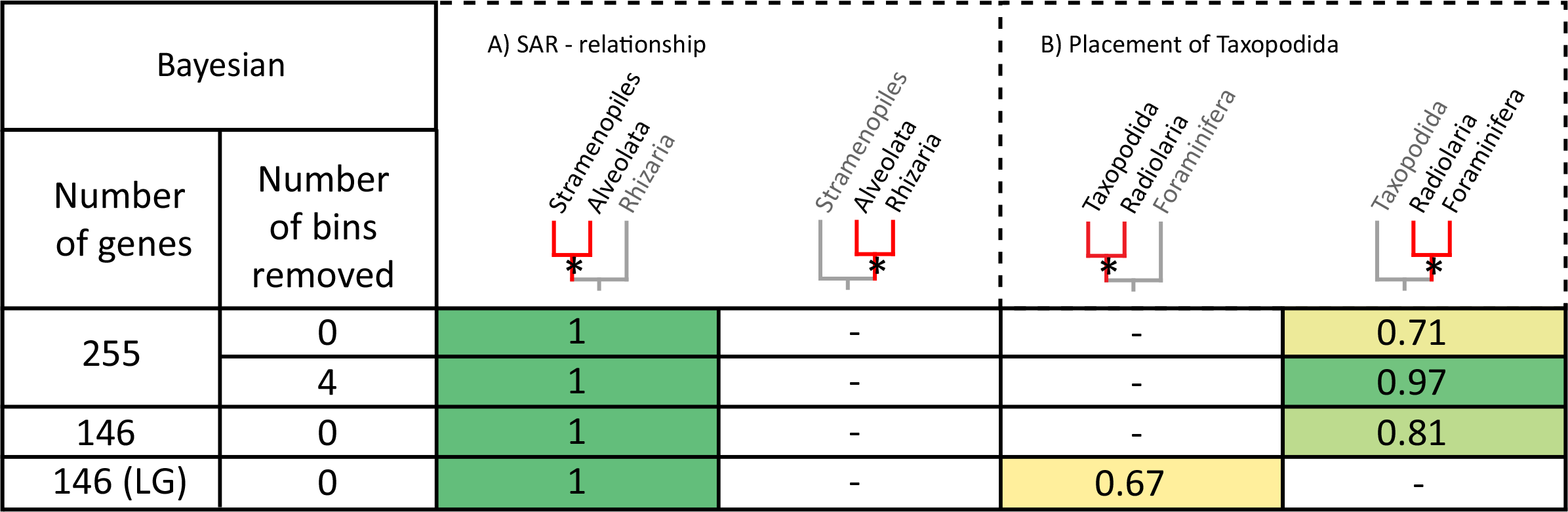
Influence of number of genes, and fast evolving sites on the Bayesian analysis using the CATGTR model, with the exception of the dataset 146 (LG) where the LG model was used. See text for discussion. **Number of genes** is the number of genes used in the concatenated data set. **Number of bins removed** equals the number of bins of fast evolving sites removed by TIGER (Cummins & McInerney 2011). The numbers in the matrix represent posterior probability for the branch marked with an asterisk. (A) The internal relationship in SAR, the first tree represents the monophyly of Stramenopiles and Alveolates, with Rhizaria as sister. The second tree represents the monophyly of Alveolates and Rhizaria, with Stramenopiles as sister. (B) The placement of Taxopodida: The first tree is the support value for Taxopodida as sister to Radiolarians, with Foraminifera as outgroup. In the second tree Taxopodida is basal in Retaria with the values showing support for the monophyly of Radiolaria and Foraminifera, excluding Taxopodida.

**Figure 3 – figure supplement 1:**
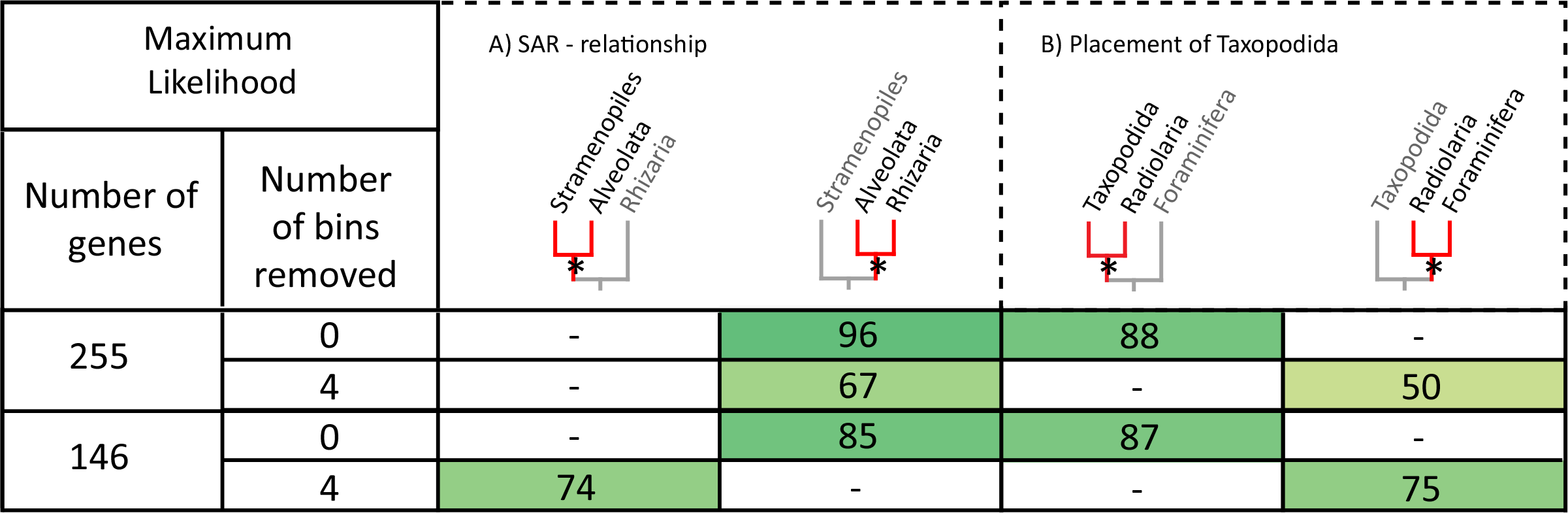
Influence of number of genes, and fast evolving sites on the ML analysis. **Number of genes** is the number of genes used in the concatenated data set. **Number of bins removed** equals the number of bins of fast evolving sites removed by TIGER (Cummins & McInerney 2011). The numbers in the matrix represent ML bootstrap values for the branch marked with an asterisk. (A) The internal relationship in SAR, the first tree represents the monophyly of Stramenopiles and Alveolates, with Rhizaria as sister. The second tree represents the monophyly of Alveolates and Rhizaria, with Stramenopiles as sister. (B) The placement of Taxopodida: The first tree is the support value for Taxopodida as sister to Radiolarians, with Foraminifera as outgroup. In the second tree Taxopodida is basal in Retaria with the values showing support for the monophyly of Radiolaria and Foraminifera, excluding Taxopodida.

**Figure 9-figure supplement 1:**
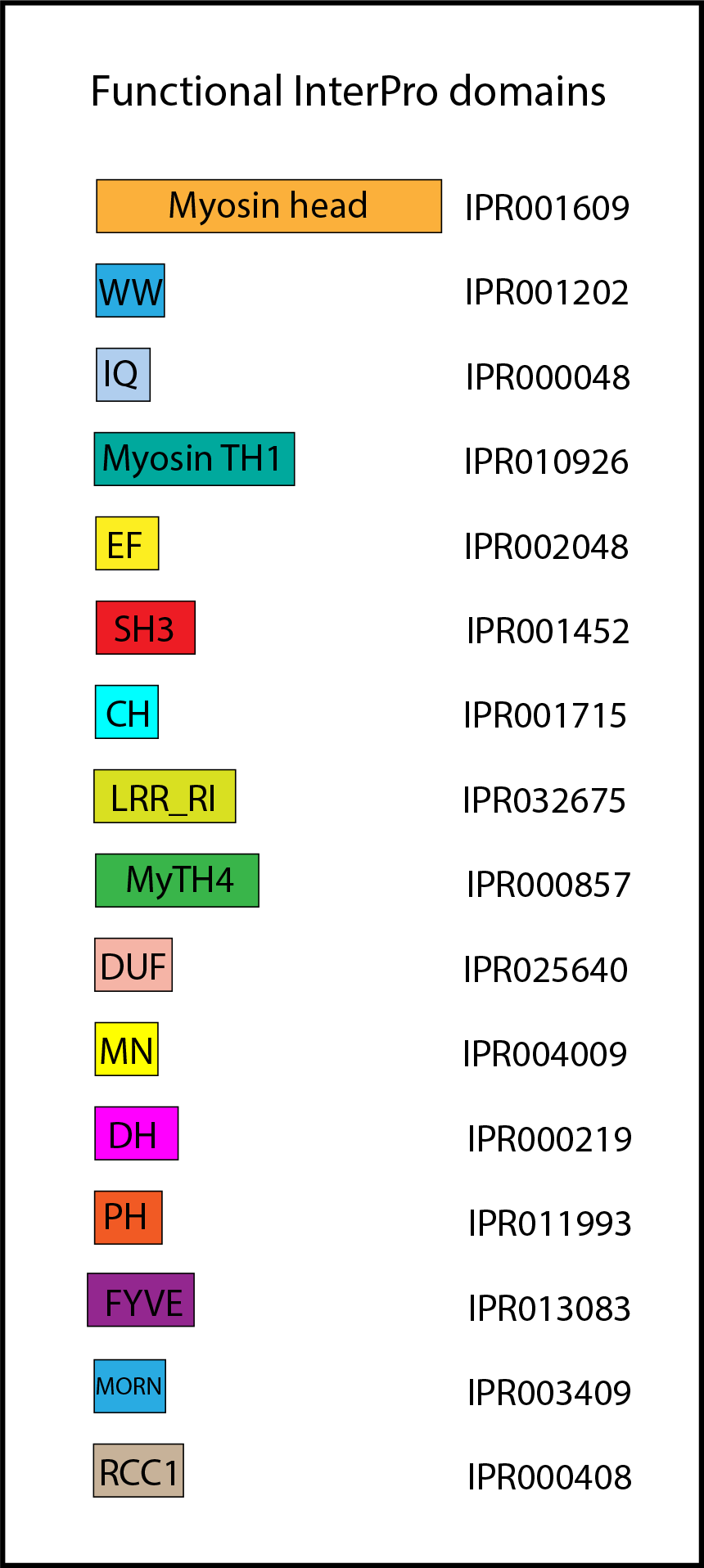
InterPro domains of myosins annotated with InterProscan (Jones et al. 2014, Mitchell et al. 2015).

**Figure 9-figure supplement 2:**
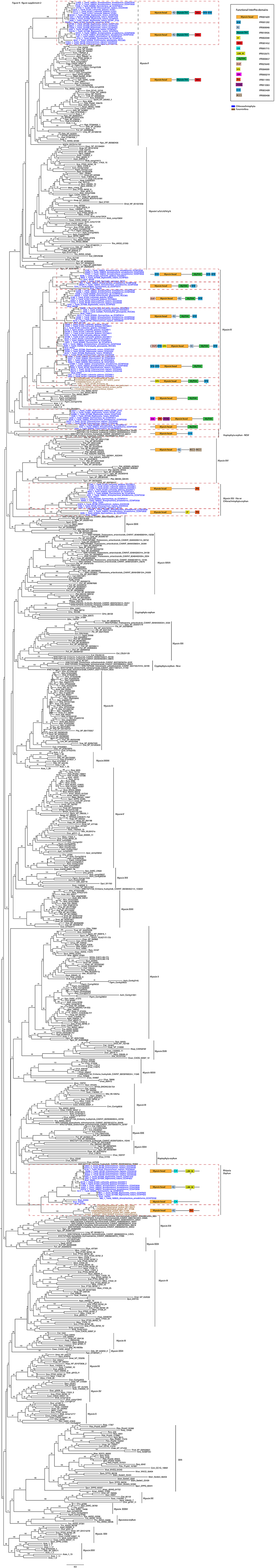
Maximum likelihood tree of myosin showing all branches. Names of taxa and myosin classes are from of Sebé-Pedrós et al. (2014), except newly discovered sequences from MMETSP and new myosin classes. The scale bar equals the mean number of substitutions per site

**Supplementary table S1:**
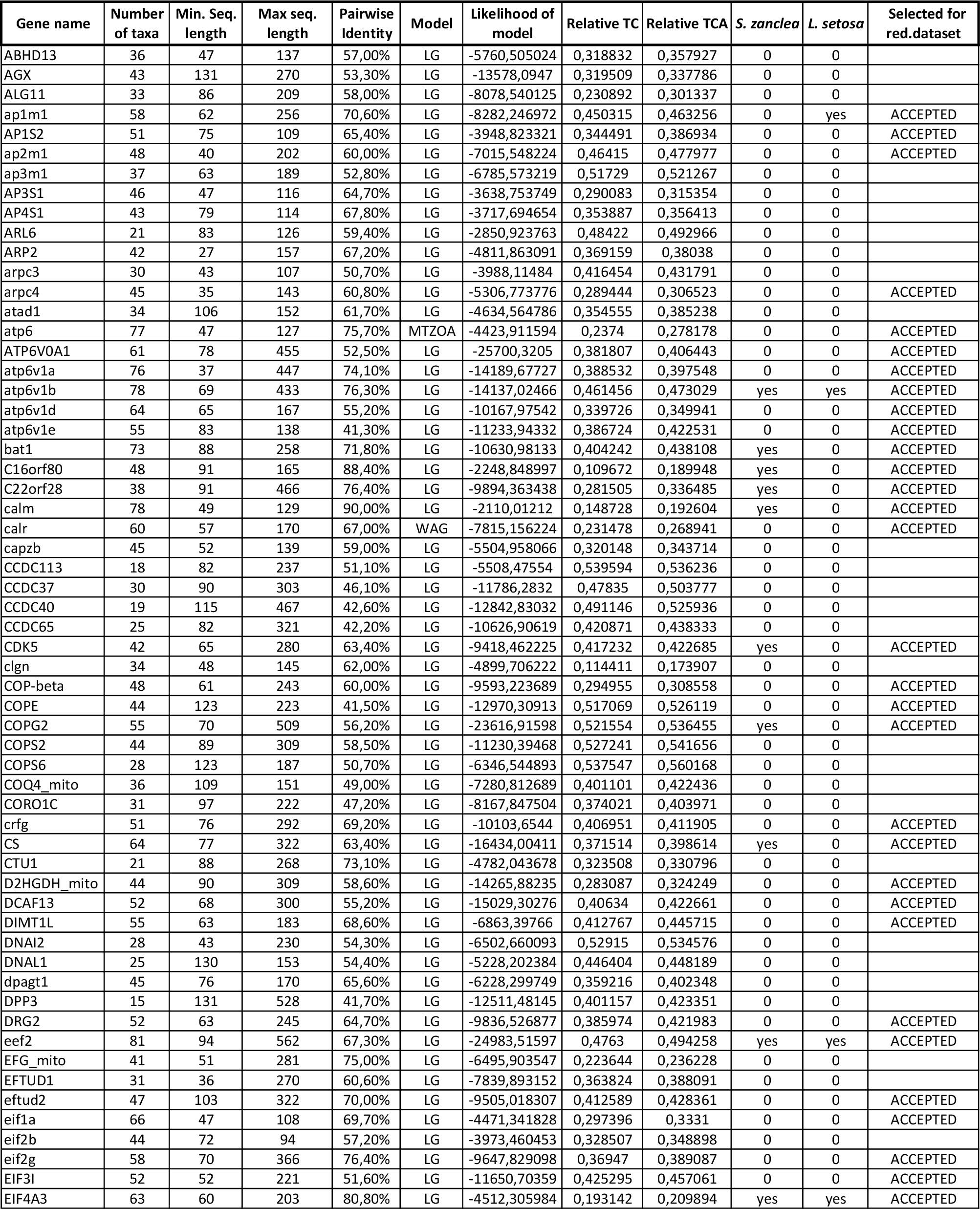
Statistics for single gene alignments. **Gene name**: Name of the gene alignment, adopted from Burki et. al (2012). **Number of taxa**: The number of taxa in the gene alignment after BIR and selection in SCaFos. **Min seq. length**: The lenght of the shortest sequence in the alignment in AA. **Max seq. length**: Length of the longest sequence in the alignment in AA. **Pairwise Identity**: Identity calculated for all pairs of sequneces in the alignment. **Model**: The model chosen as the best fitting the sequence data in the alignment by RAxML (Stamatakis 2014). **Likelihood of model**:The likelihood of the model chosen by RAxML. **Realtive TC and Realtive TCA**: Relative Tree Certainity for the phylogentic tree of the gene calculate from the boostrap trees (Salichos et al. 2014). **S. zanclea**: indicaties if the newly sequenced Sticholonche zanclea is present in the gene alignment L. **setosa**: indicaties if the newly sequenced Lithomelissa setosa is present in the gene alignment. **Selected for reduced dataset**: about half of the genes were selected for a smaller dataset. Genes marked with ACCEPT was selected to the 147-genes dataset.

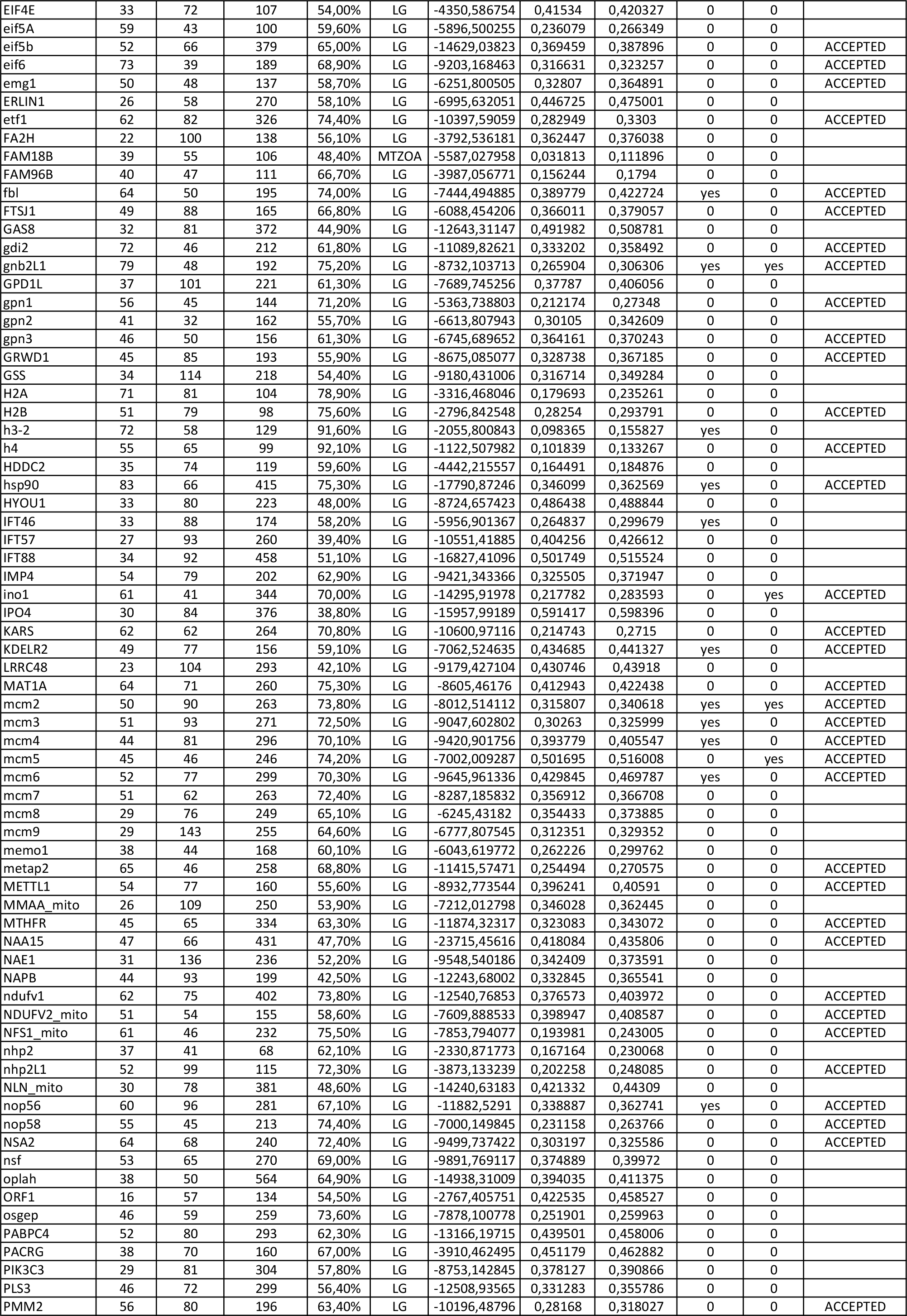

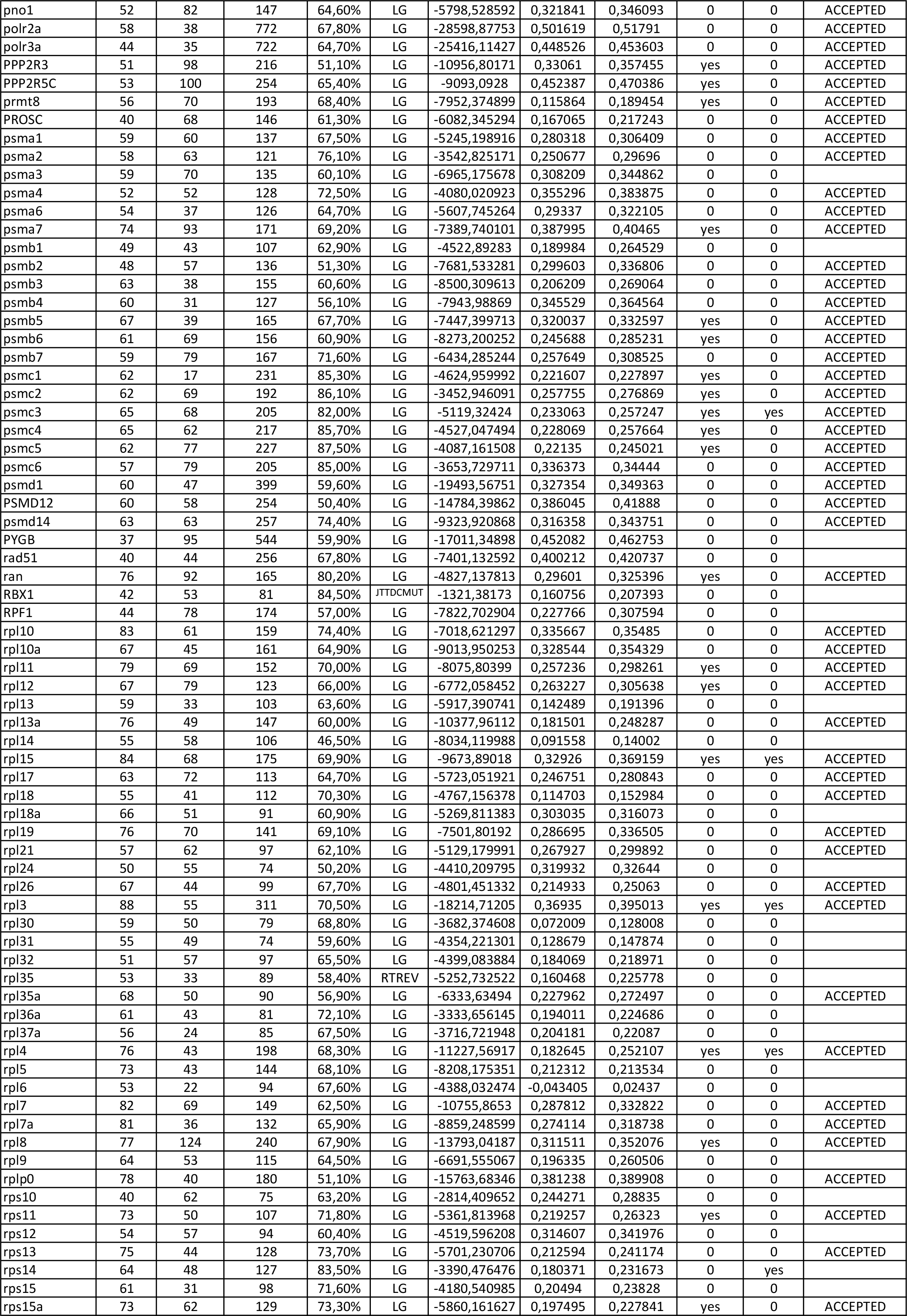

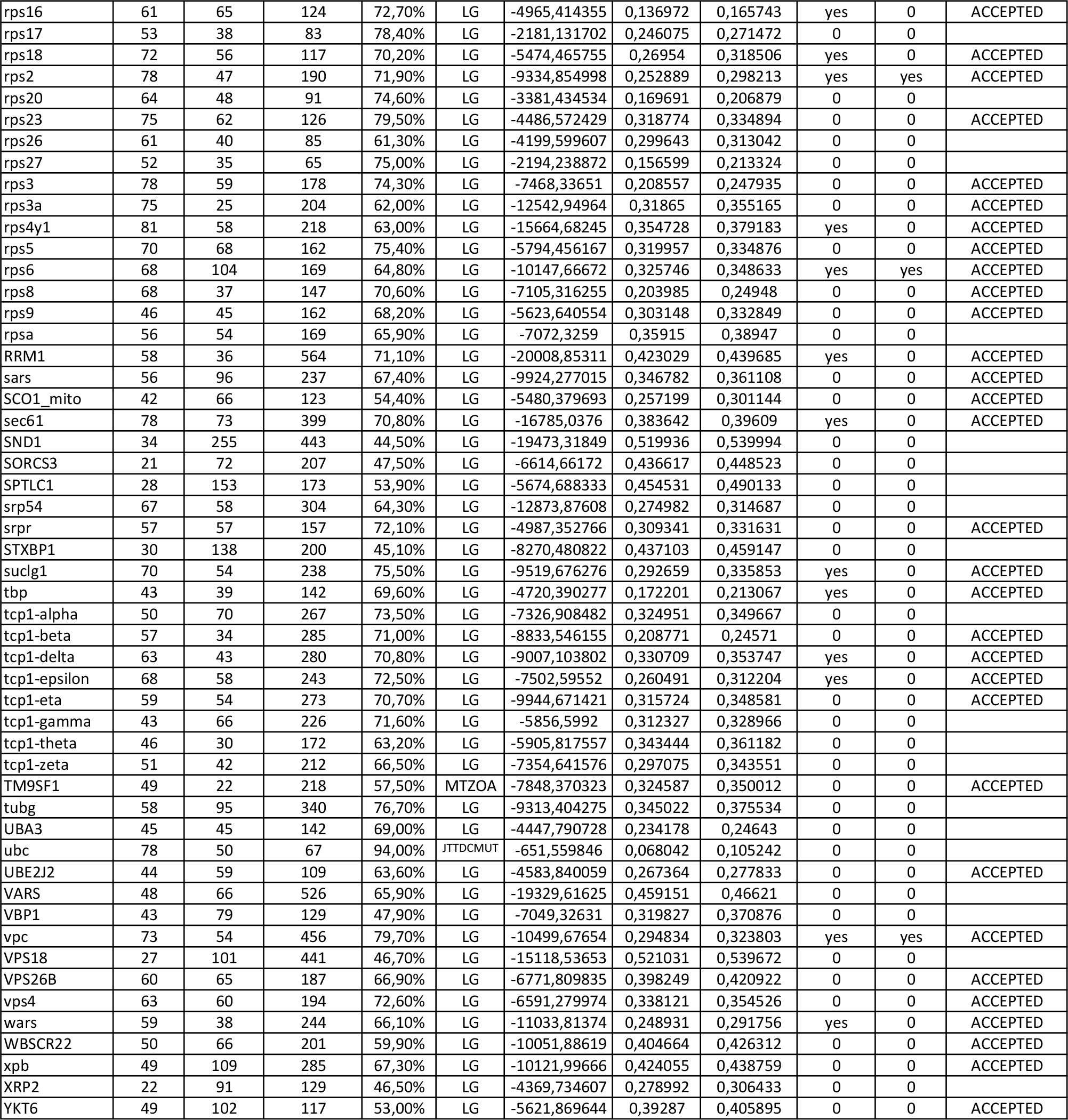

**Supplementary table S2:**
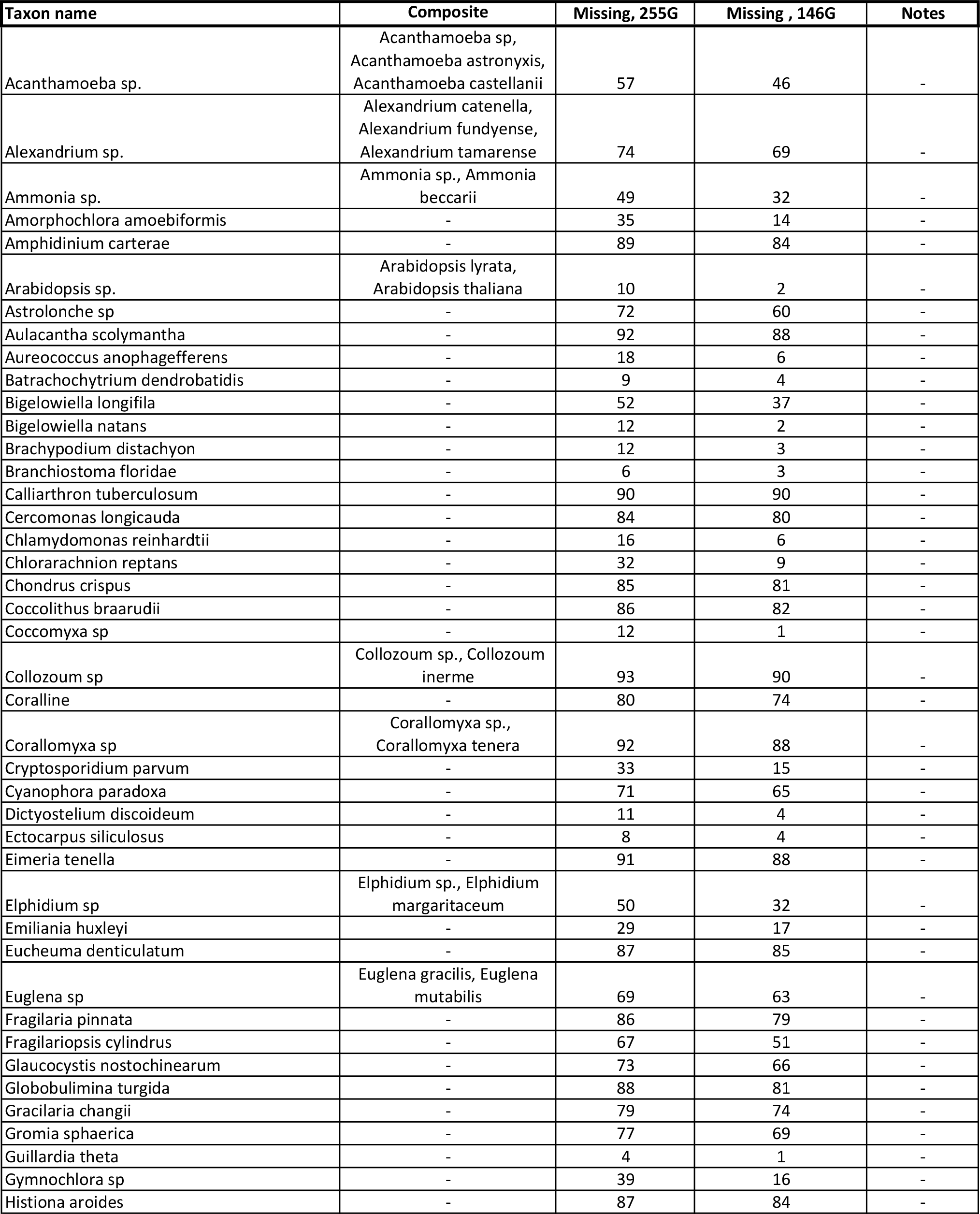
Statistics for the taxa in the multigene datasets. **Taxon name**: the name of the taxon as used in the multigene trees and alignmnent. **Composite**: Some closely related spiecies have been merged into one sequence in the alignment. **Missing, 255G**: The percentage of missing data for the given taxon in the 255 gene alignment. **Missing, 146G**: The percentage of missing data for the given taxon in the 255 gene alignment. **Notes**: Taxa marked EPA have been placed with the Evolutinary Placement Algorithm (Berger, et al. 2011), taxa marked with RogueNaRok was identified as jumping taxa and removed before concatination (Aberer et al., 2013)

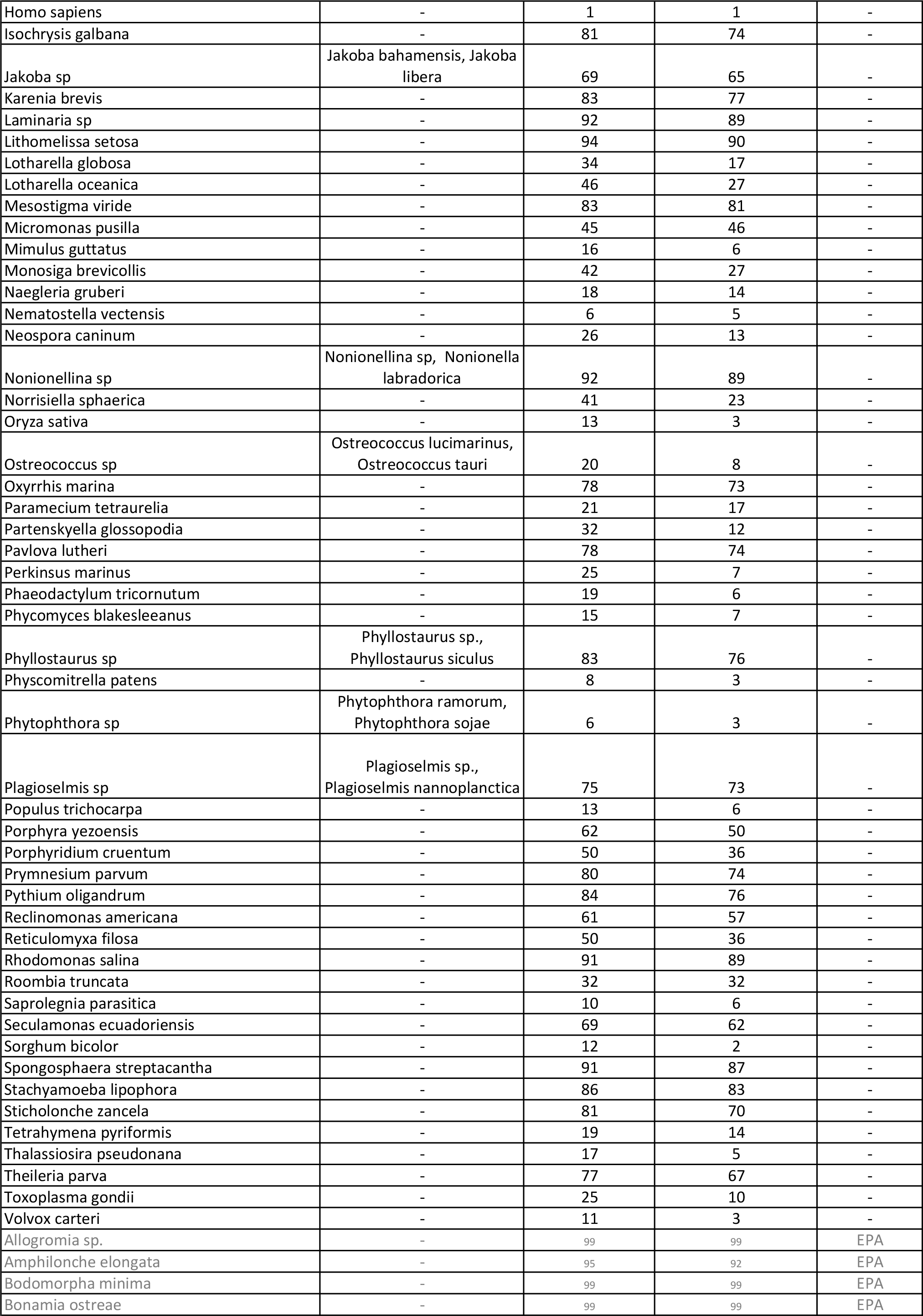

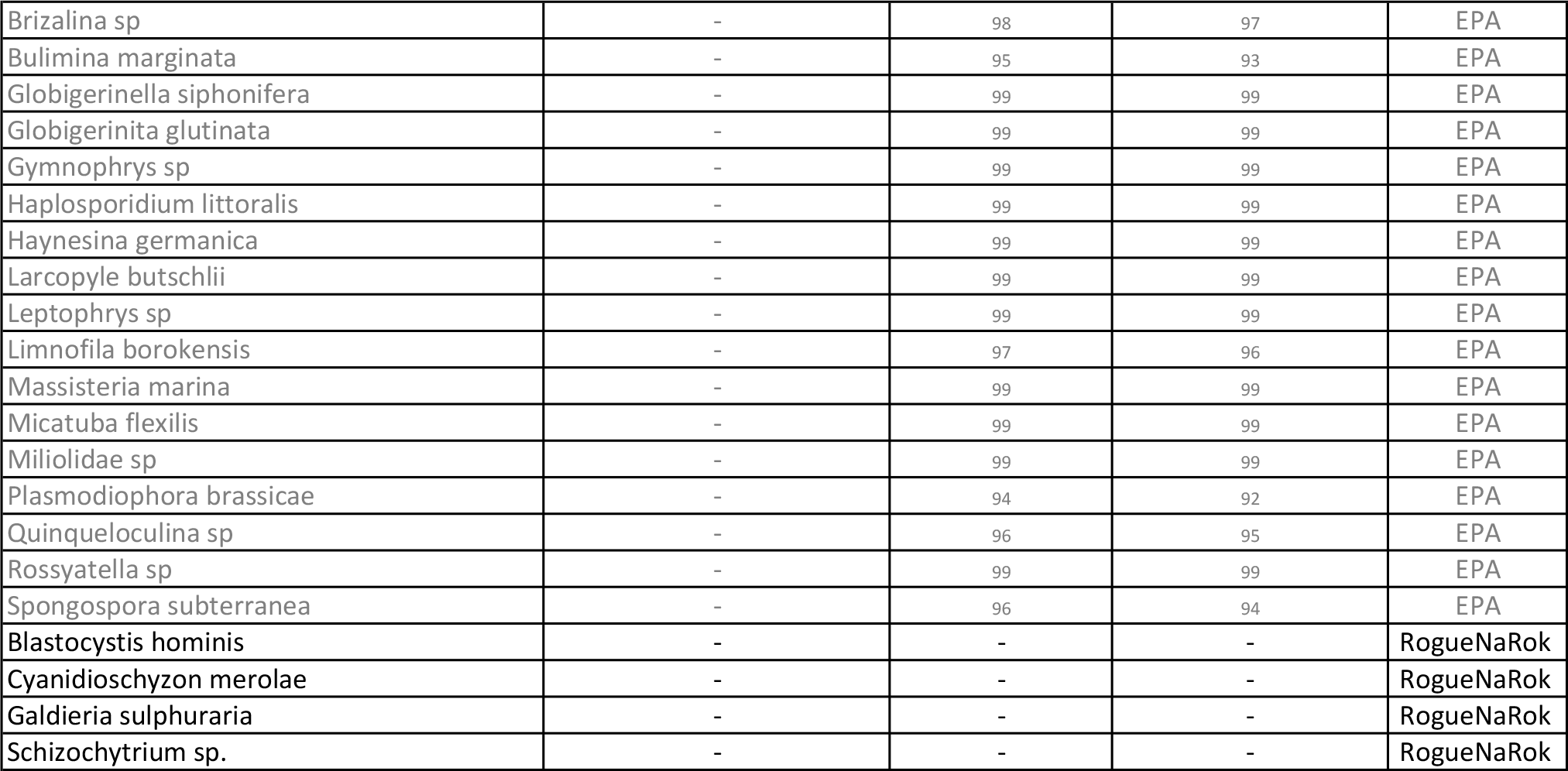

**Supplementary table S3:**
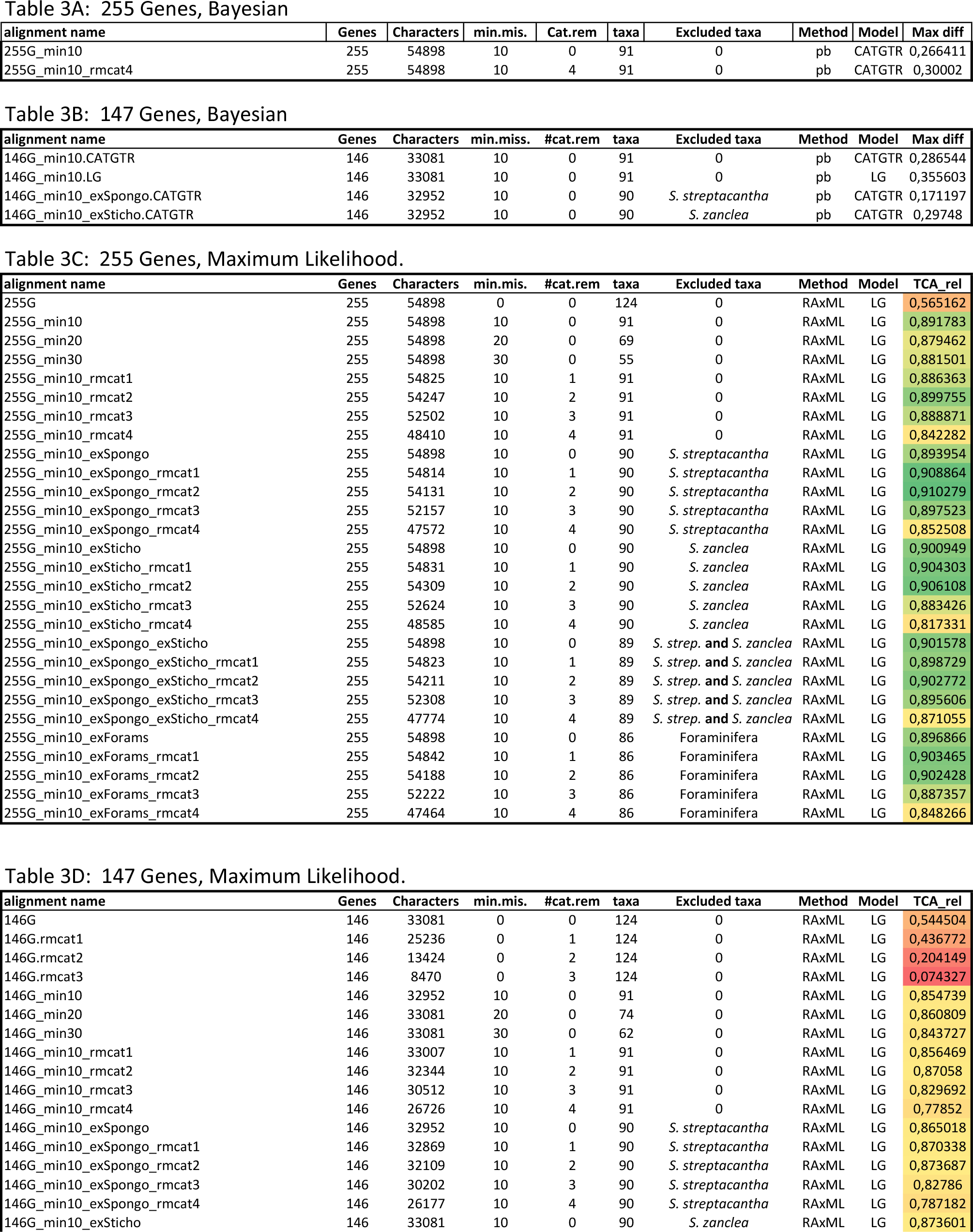
Statistics for the concatenated alignments. **Genes**: the number of genes in the alignment. Characters: The number of amino acids in the alignment. **Min.mis**: the lowest allowed percentage for missing data for a taxon in the alingnment. Cat.rem.: the number of categories of fast evolving sites removed. **Taxa**: the number of taxa in the alignment. **Excluded taxa**: The name of excluded taxa, if any. Method: The method used for phylogenetic inference pb=bayesian inference with Phylobayes mpi version 1.5a (Lartillot et al., 2013), RAxML= maximum likelihood inference with RAxML v 8.0.26 (Stamatkis 2014). Model: The model used for the inference. **Max.diff**: (only for Bayesian), maximum difference in posterior probability supportbetween two chains that ran independenlty. **TCA_rel**: (only for maximum likelihood),The relative Tree certainity index for all nodes (Salichos et al., 2014), calculated on the bootstrap files for the inferrence.

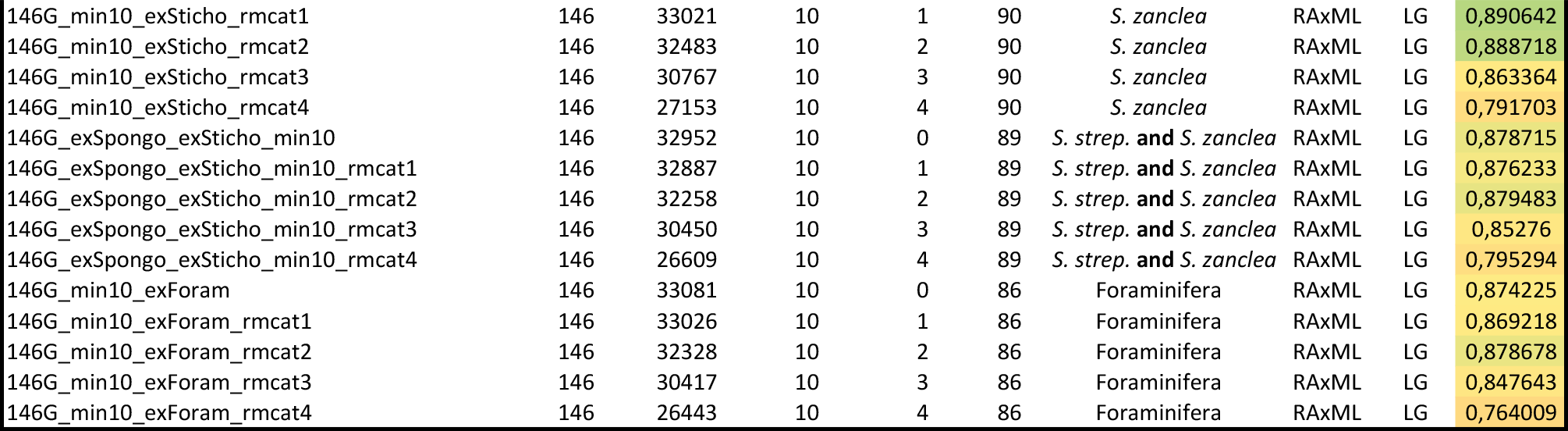

**Supplementary table S4:**
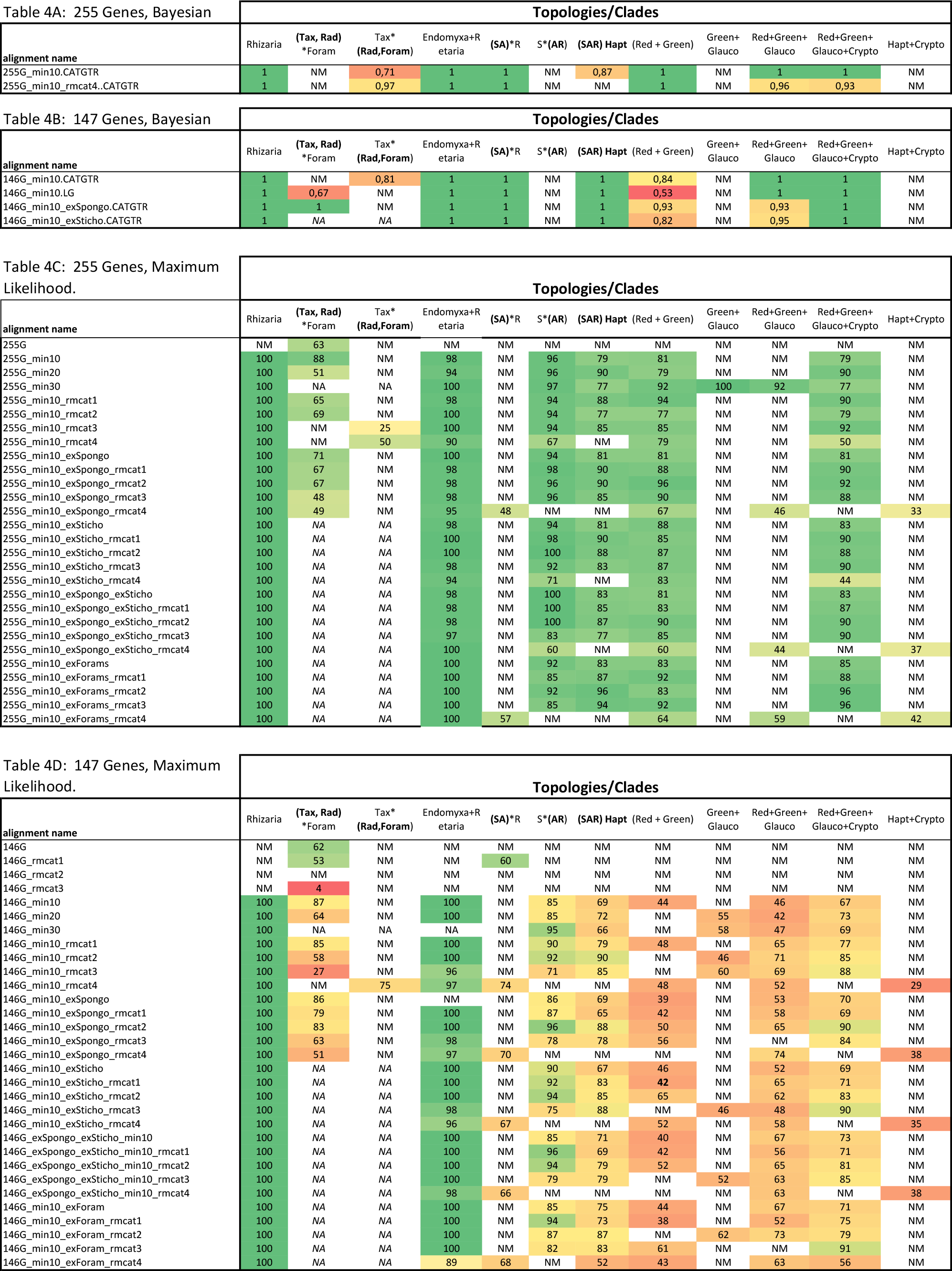
The posterior probability (bayesian) and bootstrap support (maximum likelihood) for selected nodes. The support values has been heatmapped with a three color scheme, Green color= high support, yellow=medium support, red= low support. NM= not monophyletic, i.e. the clade or node does not exist for that particular inference. NA=Not applicable, i.e. the clade or node does not exist because some taxa has been removed. For the specifics for each alignment refer to table S3. Tax=Taxopodida; Rad=Radiolaria; Foram=Foraminifer; SAR=Stramenopile, Alveolates, Rhizaria; Red=Rhodophyta; Green=Chlorophytes; Glauco=Glaucophyta; Crypto=Cryptophyta

## References

Aberer AJ, Krompass D, Stamatakis A (2013) Pruning rogue taxa improves phylogenetic accuracy: An efficient algorithm and webservice. Syst Biol 62:162–166

Anderson OR (1976a) Fine structure of a collodarian radiolarian (Sphaerozoum punctatum Muller 1858) and cytoplasmic changes during reproduction. Mar Micropaleontol 1:287–297

Anderson OR (1976b) Ultrastructure of a colonial radiolarian collozoum inerme and a cytochemical determination of the role of its zooxanthellae. Tissue Cell 8:195–208

Anderson OR (1978) Light and electron microscopic observations of feeding behavior, nutrition, and reproduction in laboratory cultures of Thalassicolla nucleata. Tissue Cell 10:401–12

Anderson OR (1983) Radiolaria, 2nd edn. Springer-Verlag, New York, USA

Ashkenazy H, Erez E, Martz E, Pupko T, Ben-Tal N (2010) ConSurf 2010: Calculating evolutionary conservation in sequence and structure of proteins and nucleic acids. Nucleic Acids Res 38

Balzano S, Corre E, Decelle J, Sierra R, Wincker P, Silva C, Poulain J, Pawlowski J, Not F (2015) Transcriptome analyses to investigate symbiotic relationships between marine protists. Front Microbiol 6:98: 1–14

Bass D, Chao EE-Y, Nikolaev S, Yabuki A, Ishida K-I, Berney C, Pakzad U, Wylezich C, Cavalier-Smith T (2009) Phylogeny of novel naked Filose and Reticulose Cercozoa: Granofilosea cl. n. and Proteomyxidea revised. Protist 160:75109

Berger S a., Krompass D, Stamatakis,A (2011) Performance, accuracy, and Web server for evolutionary placement of short sequence reads under maximum likelihood. Syst Biol 60:291–302

Berger SA, Stamatakis,A (2011) Aligning short reads to reference alignments and trees. Bioinformatics 27:2068–75

Boczkowska M, Rebowski G, Kast DJ, Dominguez,R (2014) Structural analysis of the transitional state of Arp2/3 complex activation by two actin-bound WCAs. Nat Commun 5:3308

Boczkowska M, Rebowski G, Petoukhov M V., Hayes DB, Svergun DI, Dominguez,R (2008) X-ray scattering study of activated Arp2/3 complex with bound actin-WCA. Structure 16:695–704

Bolger AM, Lohse M, Usadel,B (2014) Trimmomatic: A flexible trimmer for Illumina sequence data. Bioinformatics 30:2114–2120

Bowser, SS (2002) Reticulopodia: Structural and Behavioral Basis for the Suprageneric Placement of Granuloreticulosan Protists. J Foraminifer Res 32:440–447

Brouhard GJ, Rice,LM (2014) The contribution of αβ-tubulin curvature to microtubule dynamics. J Cell Biol 207:323–334

Brown MW, Kolisko M, Silberman JD, Roger,AJ (2012) Aggregative multicellularity evolved independently in the eukaryotic supergroup Rhizaria. Curr Biol 22:1123–1127

Burki F, Corradi N, Sierra R, Pawlowski J, Meyer GR, Abbott CL, Keeling,PJ (2013) Phylogenomics of the intracellular parasite Mikrocytos mackini reveals evidence for a mitosome in Rhizaria. Curr Biol 23:1–7

Burki F,Kaplan M,Tikhonenkov D V,Zlatogursky V,Minh BQ,Radaykina L V,Smirnov A,Mylnikov P,Keeling PJ,Keeling,PJ (2016) Untangling the early diversification of eukaryotes : a phylogenomic study of the evolutionary origins of Centrohelida, Haptophyta and Cryptista. :1–10

Burki F,Keeling PJ (2014) Rhizaria. Curr Biol 24:R103–R107

Burki F,Kudryavtsev A,Matz M V,Aglyamova G V,Bulman S,Fiers M,Keeling PJ, Pawlowski,J (2010) Evolution of Rhizaria: new insights from phylogenomic analysis of uncultivated protists. BMC Evol Biol 10:377

Burki F,Okamoto N,Pombert J-F,Keeling PJ (2012) The evolutionary history of haptophytes and cryptophytes: phylogenomic evidence for separate origins. Proc R Soc B Biol Sci 279:2246–2254

Burki F,Shalchian-Tabrizi K,Minge M,Skjaeveland A,Nikolaev SI,Jakobsen KS,Pawlowski J (2007) Phylogenomics reshuffles the eukaryotic supergroups. PLoS One 2:e790

Burki F,Shalchian-Tabrizi K, Pawlowski,J (2008) Phylogenomics reveals a new “megagroup” including most photosynthetic eukaryotes. Biol Lett 4:366–9

Cachon J,Cachon M (1971) Le system axopodial des Radiolaires Nassellaires. Arch für Protistenkd 113:80–97

Cachon J,Cachon M,Febvre-Chevalier C,Febvre J (1973) Determinisme de l'edification des systemes microtubulaires stereoplasmiques d'Actinopodes. Arch für Protistenkd 115:137–153

Cachon J,Cachon M,Tilney LG,Tilney MS (1977) Movement generated by interactions between the dense material at the ends of microtubules and non-actin-containing microfilaments in Sticholonche zanclea. J Cell Biol 72:314–338

Cavalier-Smith T (1993) Kingdom Protozoa and its 18 phyla. Microbiol Rev 57:953–994

Cavalier-Smith T (2002) The phagotrophic origin of eukaryotes and phylogenetic classification of Protozoa. Int J Syst Evol Microbiol 52:297–354

Cavalier-Smith T (2009) Megaphylogeny, cell body plans, adaptive zones: causes and timing of eukaryote basal radiations. J Eukaryot Microbiol 56:26–33

Cavalier-Smith T,Chao EE,Lewis R (2015) Multiple origins of Heliozoa from flagellate ancestors: New cryptist subphylum Corbihelia, superclass Corbistoma, and monophyly of Haptista, Cryptista, Hacrobia and Chromista. Mol Phylogenet Evol 93:331–362

Cavalier-Smith T,Guy L,Saw JH,Ettema TJG,Eme L,Sharpe SC,Brown MW,Irimia M,Roy SW (2014) The neomuran revolution and phagotrophic origin of eukaryotes and cilia in the light of intracellular coevolution and a revised Tree of Life. Cold Spring Harb Perspect Biol 6:a016006–a016006

Celniker G,Nimrod G,Ashkenazy H,Glaser F,Martz E,Mayrose I,Pupko T,Ben-Tal N (2013) ConSurf: Using evolutionary data to raise testable hypotheses about protein function. Isr J Chem 53:199–206

Cummins C a,McInerney JO (2011) A method for inferring the rate of evolution of homologous characters that can potentially improve phylogenetic inference, resolve deep divergence and correct systematic biases. Syst Biol 60:833–844

Dustin P (1984) Microtubules, 2nd edn. Springer-Verlag, Berlin

Febvre J (1981) The myoneme of the Acantharia (Protozoa): A new model of cellular motility. Biosystems 14:327–336

Finn RD,Bateman A,Clements J,Coggill P,Eberhardt RY,Eddy SR,Heger A,Hetherington K,Holm L,Mistry J,Sonnhammer ELL,Tate J,Punta M (2014) Pfam: the protein families database. Nucleic Acids Res 42:D222–30

Giannone G,Dubin-Thaler BJ,Rossier O,Cai Y,Chaga O,Jiang G,Beaver W,Döbereiner H-G,Freund Y,Borisy G,Sheetz MP (2007) Lamellipodial actin mechanically links myosin activity with adhesion-site formation. Cell 128:561–75

Goldstein LS (2001) Molecular motors: from one motor many tails to one motor many tales. Trends Cell Biol 11:477–82

Goley ED,Welch MD (2006) The ARP2/3 complex: an actin nucleator comes of age. Nat Rev Mol Cell Biol 7:713–726

Grain J (1986) The cytoskeleton in protists: nature, structure, and functions. Int Rev Cytol 104:153–249

Haas BJ,Papanicolaou A,Yassour M,Grabherr M,Blood PD,Bowden J,Couger MB,Eccles D,Li B,Lieber M,Macmanes MD,Ott M,Orvis J,Pochet N,Strozzi F,Weeks N,Westerman R,William T,Dewey CN,Henschel R,Leduc RD,Friedman N,Regev A (2013) De novo transcript sequence reconstruction from RNA-seq using the Trinity platform for reference generation and analysis. Nat Protoc 8:1494–1512

Habura A,Wegener L,Travis JL,Bowser SS (2005) Structural and functional implications of an unusual foraminiferal betα-tubulin. Mol Biol Evol 22:2000–2009

Hammer J a.,Wu XS (2002) Rabs grab motors: Defining the connections between Rab GTPases and motor proteins. Curr Opin Cell Biol 14:69–75

He D,Sierra R,Pawlowski J,Baldauf SL (2016) Reducing long-branch effects in multi-protein data uncovers a close relationship between Alveolata and Rhizaria. Mol Phylogenet Evol

Hou Y,Sierra R,Bassen D,Banavali NK,Habura A,Pawlowski J,Bowser SS (2013) Molecular evidence for β-tubulin neofunctionalization in Retaria (Foraminifera and Radiolarians). Mol Biol Evol 30:2487–2493

Ishida KI,Yabuki A,Ota S (2011) Research note: Amorphochlora amoebiformis gen. et comb. nov. (Chlorarachniophyceae). Phycol Res 59:52–53

Ishitani Y,Ishikawa S,Inagaki Y,Tsuchiya M,Takishita K (2011) Multigene phylogenetic analyses including diverse radiolarian species support the “Retaria” hypothesis-the sister relationship of Radiolaria and Foraminifera. Mar Micropaleontol 81:32–42

Jékely G (ed) (2007) Eukaryotic Membranes and Cytoskeleton. Springer New York, New York, NY

Jones P,Binns D,Chang HY,Fraser M,Li W,McAnulla C,McWilliam H,Maslen J,Mitchell A,Nuka G,Pesseat S,Quinn AF,Sangrador-Vegas A,Scheremetjew M,Yong SY,Lopez R, Hunter S (2014) InterProScan 5: Genome-scale protein function classification. Bioinformatics 30:1236–1240

Karcher RL,Deacon SW,Gelfand VI (2002) Motor-cargo interactions: The key to transport specificity. Trends Cell Biol 12:21–27

Kast DJ,Zajac AL,Holzbaur ELF,Ostap EM,Dominguez Correspondence R,Dominguez R (2015) WHAMM Directs the Arp2/3 Complex to the ER for Autophagosome Biogenesis through an Actin Comet Tail Mechanism. Curr Biol25:1791–1797

Katz L a (2012) Origin and Diversification of Eukaryotes. Annu Rev Microbiol:411–;427

Katz LA,Grant JR (2014) Taxon-rich phylogenomic analyses resolve the eukaryotic Tree of Life and reveal the power of subsampling by sites. Syst Biol 64:406–415

Kearse M, Moir R, Wilson A, Stones-Havas S, Cheung M, Sturrock S, Buxton S, Cooper A, Markowitz S, Duran C, Thierer T, Ashton B, Meintjes P, Drummond A (2012) Geneious Basic: An integrated and extendable desktop software platform for the organization and analysis of sequence data. Bioinformatics 28:1647–1649

Keeling PJ, Burki F, Wilcox HM, Allam B, Allen EE, Amaral Zettler L, Armbrust EV, Archibald JM, Bharti AK, Bell CJ, Beszteri B, Bidle KD, Cameron CT, Campbell L, Caron D a., Cattolico RA, Collier JL, Coyne K, Davy SK, Deschamps P, Dyhrman ST, Edvardsen B, Gates RD, Gobler CJ, Greenwood SJ, Guida SM, Jacobi JL, Jakobsen KS, James ER, Jenkins B, John U, Johnson MD, Juhl AR, Kamp A, Katz L a., Kiene R, Kudryavtsev A, Leander BS, Lin S, Lovejoy C, Lynn D, Marchetti A, McManus G, Nedelcu AM, Menden-Deuer S, Miceli C, Mock T, Montresor M, Moran MA, Murray S, Nadathur G, Nagai S, Ngam PB, Palenik B, Pawlowski J, Petroni G, Piganeau G, Posewitz MC, Rengefors K, Romano G, Rumpho ME, Rynearson T, Schilling KB, Schroeder DC, Simpson AGB, Slamovits CH, Smith DR, Smith GJ, Smith SR, Sosik HM, Stief P, Theriot E, Twary SN, Umale PE, Vaulot D, Wawrik B, Wheeler GL, Wilson WH, Xu Y, Zingone A, Worden AZ (2014) The Marine Microbial Eukaryote Transcriptome Sequencing Project (MMETSP): illuminating the functional diversity of eukaryotic life in the oceans through transcriptome sequencing (RG Roberts, Ed.). PLoS Biol 12:e1001889

Kolisko M, Boscaro V, Burki F, Lynn DH, Keeling PJ (2014) Single-cell transcriptomics for microbial eukaryotes. Curr Biol 24:R1081–R1082

Kollmar M, Lbik D, Enge S (2012) Evolution of the eukaryotic ARP2/3 activators of the WASP family: WASP, WAVE, WASH, and WHAMM, and the proposed new family members WAWH and WAML. BMC Res Notes 5:88

Krabberød AK, Bråte J, Dolven JK, Ose RF, Klaveness D, Kristensen T, Bjørklund KR, Shalchian-Tabrizi K (2011) Radiolaria Divided into Polycystina and Spasmaria in Combined 18S and 28S rDNA Phylogeny. PLoS One 6:e23526

Kumar S, Krabberød AK, Neumann RS, Michalickova K, Zhang X, Zhao S, Shalchian-Tabrizi K (2015) BIR Pipeline for Preparation of Phylogenomic Data. Evol Bioinforma 11:79–83

Lartillot N, Philippe H (2004) A Bayesian mixture model for across-site heterogeneities in the amino-acid replacement process. Mol Biol Evol 21:1095–1109

Lartillot N, Rodrigue N, Stubbs D, Richer J (2013) Phylobayes mpi: Phylogenetic reconstruction with infinite mixtures of profiles in a parallel environment. Syst Biol 62:611–615

Lee JJ, Anderson OR (Eds) (1991) Biology of Foraminifera. Academic Press, London, UK, London

Liang J, Cai W, Sun Z (2014) Single-Cell sequencing technologies: current and future. J Genet Genomics 41:513–528

Liu N, Liu L, Pan X (2014) Single-cell analysis of the transcriptome and its application in the characterization of stem cells and early embryos. Cell Mol Life Sci 71:2707–2715

Löwe J, Li H, Downing KH, Nogales E (2001) Refined structure of alpha betα-tubulin at 3.5 A resolution. J Mol Biol

Margulis L (1990) Handbook of protoctista : the structure, cultivation, habitats, and life histories of the eukaryotic microorganisms and their descendants exclusive of animals, plants and fungi : a guide to the algae, ciliates, foraminifera, sporozoa, water molds, slime m (L Margulis, Ed.). Jones and Bartlett Publishers, Boston

Mattila PK, Lappalainen P (2008) Filopodia: molecular architecture and cellular functions. Nat Rev Mol Cell Biol 9:446–454

Mitchell A, Chang HY, Daugherty L, Fraser M, Hunter S, Lopez R, McAnulla C, McMenamin C, Nuka G, Pesseat S, Sangrador-Vegas A, Scheremetjew M, Rato C, Yong SY, Bateman A, Punta M, Attwood TK, Sigrist CJA, Redaschi N, Rivoire C, Xenarios I, Kahn D, Guyot D, Bork P, Letunic I, Gough J, Oates M, Haft D, Huang H, Natale DA, Wu CH, Orengo C, Sillitoe I, Mi H, Thomas PD, Finn RD (2015) The InterPro protein families database: The classification resource after 15 years. Nucleic Acids Res 43:D213–D221

Mogilner A, Keren K (2009) The Shape of Motile Cells. Curr Biol 19:R762–R771

Moreira D, Heyden S von der, Bass D, Löpez-Garía P, Chao E, Cavalier-Smith T (2007) Global eukaryote phylogeny: Combined small-and large-subunit ribosomal DNA trees support monophyly of Rhizaria, Retaria and Excavata. Mol Phylogenet Evol 44:255–266

Nikolaev SI, Berney C, Fahrni JF, Bolivar I, Polet S, Mylnikov AP, Aleshin V V, Petrov NB, Pawlowski J (2004) The twilight of Heliozoa and rise of Rhizaria, an emerging supergroup of amoeboid eukaryotes. Proc Natl Acad Sci U S A 101:8066–8071

Parfrey LW, Grant J, Tekle YI, Lasek-Nesselquist E, Morrison HG, Sogin ML, Patterson DJ, Katz L a (2010) Broadly sampled multigene analyses yield a well-resolved eukaryotic tree of life. Syst Biol 59:518–533

Pattengale ND, Swenson KM, Moret BME (2010) Uncovering hidden phylogenetic consensus. In: Lecture Notes in Computer Science (including subseries Lecture Notes in Artificial Intelligence and Lecture Notes in Bioinformatics).p 128–139

Pawlowski J (2008) The twilight of Sarcodina: a molecular perspective on the polyphyletic origin of amoeboid protists. Protistology 5:281–302

Pawlowski J, Burki F (2009) Untangling the phylogeny of amoeboid protists. J Eukaryot Microbiol 56:16–25

Philippe H, Zhou Y, Brinkmann H, Rodrigue N, Delsuc F (2005) Heterotachy and long-branch attraction in phylogenetics. BMC Evol Biol 5:50

Picelli S, Faridani OR, Björklund AK, Winberg G, Sagasser S, Sandberg R (2014) Full-length RNA-seq from single cells using Smart-seq2. Nat Protoc 9:171–181

Pollard TD (2007) Regulation of actin filament assembly by Arp2/3 complex and formins. Annu Rev Biophys Biomol Struct 36:451–477

Richards T a, Cavalier-Smith T (2005) Myosin domain evolution and the primary divergence of eukaryotes. Nature 436:1113–8

Rojas AM, Fuentes G, Rausell A, Valencia A (2012) The Ras protein superfamily: Evolutionary tree and role of conserved amino acids. J Cell Biol 196:189–201

Rouiller I, Xu XP, Amann KJ, Egile C, Nickell S, Nicastro D, Li R, Pollard TD, Volkmann N, Hanein D (2008) The structural basis of actin filament branching by the Arp2/3 complex. J Cell Biol 180:887–895

Roure B, Rodriguez-Ezpeleta N, Philippe H (2007) SCaFoS: a tool for selection, concatenation and fusion of sequences for phylogenomics. BMC Evol Biol 7 Suppl 1:S2

Salichos L, Stamatakis A, Rokas A (2014) Novel information theory-based measures for quantifying incongruence among phylogenetic trees. Mol Biol Evol 31:1261–1271

Schliwa M, Woehlke G (2003) Molecular motors. Nature 422:759–765

Seabra MC, Coudrier E (2004) Rab GTPases and myosin motors in organelle motility. Traffic 5:393–9

Sebé-Pedrós A, Grau-Bové X, Richards TA, Ruiz-Trillo I (2014) Evolution and classification of myosins, a paneukaryotic whole-genome approach. Genome Biol Evol 6:290–305

Sierra R, Cañas-Duarte SJ, Burki F, Schwelm A, Fogelqvist J, Dixelius C, González-García LN, Gile GH, Slamovits CH, Klopp C, Restrepo S, Arzul I, Pawlowski J (2015) Evolutionary origins of rhizarian parasites. Mol Biol Evol 33:msv340–

Sierra R, Matz M V., Aglyamova G, Pillet L, Decelle J, Not F, Vargas C de, Pawlowski J (2013) Deep relationships of Rhizaria revealed by phylogenomics: A farewell to Haeckel's Radiolaria. Mol Phylogenet Evol 67:53–59

Stamatakis A (2014) RAxML version 8: A tool for phylogenetic analysis and post-analysis of large phylogenies. Bioinformatics 30:1312–1313

Stamatakis A, Komornik Z, Berger SA (2010) Evolutionary placement of short sequence reads on multi-core architectures. In: 2010 ACS/IEEE International Conference on Computer Systems and Applications, AICCSA 2010.p 1–44

Sugiyama K, Hori RS, Kusunoki Y, Matsuoka A (2008) Pseudopodial features and feeding behavior of living nassellarians Eucyrtidium hexagonatum Haeckel, Pterocorys zancleus (Muler) and Dictyocodon prometheus Haeckel. Paleontol Res 12:209–222

Suzuki NO, Aita YO (2011) Radiolaria: achievements and unresolved issues: taxonomy and cytology. Plankt Benthos Res 6:69–91

Townsend JP (2007) Profiling phylogenetic informativeness. Syst Biol 56:222–231

Townsend JP, Su Z, Tekle YI, Ownsend JEPT, Huo ZSU (2012) Phylogenetic signal and noise: predicting the power of a data set to resolve phylogeny. Syst Biol 61:835–49

Travis JL, Allen RD (1981) Studies on the Motility of the Foraminifera .1. Ultrastructure of the Reticulopodial Network of Allogromia-Laticollaris (Arnold). J Cell Biol 90:211–221

Travis JL, Bowser SS (1986) A new model of reticulopodial motility and shape: evidence for a microtubule-based motor and an actin skeleton. Cell Motil Cytoskeleton 6:2–14

Ura S, Pollitt AY, Veltman DM, Morrice N a., MacHesky LM, Insall RH (2012) Pseudopod growth and evolution during cell movement is controlled through SCAR/WAVE dephosphorylation. Curr Biol 22:553–561

Vale RD (2003) The molecular motor toolbox for intracellular transport. Cell 112:467–480

Volkmann N, Amann KJ, Stoilova-McPhie S, Egile C, Winter DC, Hazelwood L, Heuser JE, Li R, Pollard TD, Hanein D (2001) Structure of Arp2/3 complex in its activated state and in actin filament branch junctions. Science 293:2456–2459

Welnhofer E a, Travis JL (1998) Evidence for a direct conversion between two tubulin polymers--microtubules and helical filaments--in the foraminiferan, Allogromia laticollaris. Cell Motil Cytoskeleton 41:107–116

Wickstead B, Gull K (2011) The evolution of the cytoskeleton. J Cell Biol 194:513–525

Xu X-P, Rouiller I, Slaughter BD, Egile C, Kim E, Unruh JR, Fan X, Pollard TD, Li R, Hanein D, Volkmann N (2011) Three-dimensional reconstructions of Arp2/3 complex with bound nucleation promoting factors. EMBO J 31:236–247

Zhang J, Kobert K, Flouri T, Stamatakis A (2013) PEAR: A fast and accurate Illumina Paired-End reAd mergeR. Bioinformatics 30:1–7

